# Comprehensive molecular profiling of single-cell proteome via gel electrophoresis and 3D single-molecule imaging

**DOI:** 10.64898/2026.01.14.699423

**Authors:** Latiefa Kamarulzaman, Sooyeon Kim, Takuya Hidaka, Misaki Tsuchida, Yuichi Taniguchi

**Author notes:** (Corresponding author: Yuichi Taniguchi,).

## Abstract

Recent advances in shotgun proteomics and immunoassays have yielded powerful single-cell proteomics technologies. However, current methods lack the sensitivity required to comprehensively quantify protein abundances in individual cells. Here, we present single-cell PAGE-PISA, an ultra-sensitive proteome profiling strategy that combines gel electrophoresis with 3D single-molecule fluorescence imaging. Our approach labels all proteins in single cells with fluorescent dyes, separates them by electrophoresis, and counts with single-molecule resolution. This technique quantified over 10^7^ protein copies from a single mammalian cell with the sensitivity to detect low-abundance proteins down to 10^4^ copies per species. Single-cell PAGE-PISA successfully classified cells into distinct cell types based on their proteomic profiles. Furthermore, our single-cell proteome data strongly correlated with predicted developmental states during cardiomyocyte differentiation, providing complementary information to single-cell transcriptome data. Together, single-cell PAGE-PISA enables highly sensitive and quantitative proteome profiling at the single-cell level, capturing subtle proteomic differences that distinguish diverse cellular states.

## Introduction

Heterogeneity occurs at multiple levels of molecular biology and is inherent in many biological processes. For many years, the central dogma of molecular biology has been explored primarily through conventional bulk analyses. However, such approaches provide an average measurement (e.g., protein abundance, gene expression) across a population of cells. While bulk analyses have been useful for distinguishing between diseased and healthy tissues^1^, they often reflect dominant biological traits. To address this issue, single-cell technologies have been developed to capture the unique molecular profiles of individual cells and provide insights into cellular heterogeneity. One widely adopted approach is single-cell transcriptome profiling, which captures RNA expression levels across a broad range of transcripts at the single-cell resolution, enabling clustering of diverse cell types and states within complex populations^2,3,4^. Despite its potential, RNA expression levels cannot accurately predict protein abundance^5,6,7,8,9^. This is because protein abundance is highly dynamic and greatly influenced by various processes, such as protein degradation and post-transcriptional and translational modifications^10,11^. Therefore, it is imperative to perform direct proteome profiling at the single-cell level to comprehensively capture the protein abundance and modifications that are not reflected by RNA expression.

Currently, there are two major approaches to quantifying protein expression levels in single cells, which are mass spectrometry (MS) and antibody-based analysis^12^. So far, MS-based analysis serves as the gold standard for proteomic studies, attributable to its ability to identify, characterize, and quantify proteins with high multiplexity^13^. However, MS does not provide a comprehensive proteomic analysis for single cells due to its limitations in sensitivity. This makes accurate quantification of proteins using MS even more difficult for single cells because of its low protein abundance, between 50 and 300 pg in single mammalian cell^14^, and the fact that proteins cannot be amplified like nucleic acids.

This limitation was significantly addressed by the development of SCoPE-MS^15^, which enables the quantification of over 1,000 protein groups per single cell. Since then, many MS-based single-cell proteomics technologies have been developed to improve the detection sensitivity^16,17,18,19,20,21,22,23,24^. Yet, these methods predominantly quantified highly abundant proteins (>10^4^ copies per cell), while low abundant proteins (10^1^‒10^3^ copies per cell) remain challenging to quantify^6^.

Apart from MS-based analysis, other single-cell protein analysis techniques have also been developed based on antibody labelling^25,26,27^. In 2014, Herr et al. developed single-cell Western, which involves isolating single cells into microwells, lysing them in situ, separating proteins by electrophoresis, and immobilizing them for protein abundance analysis of target proteins using antibodies. This approach offers high-throughput analysis by enabling simultaneous assay of 1,000‒2,000 single cells in less than 4 hours. This represents an improvement in both sample throughput and measurement time compared to most MS-based single-cell proteomics approaches, albeit sensitivity remains relatively the same. Recent modifications of single-cell Western using nitrocellulose blotting and enzyme-antibody conjugates have lowered the detection limit to approximately 10^3^ molecules per protein species^28^. Despite this progress, its sensitivity is still insufficient for analyzing low-abundance proteins, which remains a key impediment of single-cell Western.

To overcome these limitations, single-molecule fluorescence microscopy offers precise detection and quantification of target molecules. Recently, our group developed a custom-built light-sheet microscope called <u>p</u>lanar illumination micro<u>s</u>cope for single-molecule imaging for <u>a</u>ll purpose (PISA)^29^, which enables 3D single-molecule imaging of all target molecules within a sub-millimeter sample depth. By placing the sample plane above the optical systems for light-sheet imaging and the two objective lenses for illumination and detection below the coverslip at a tilted angle, PISA facilitates counting the number of molecules in the entire volume of a mm-sized biological specimen.

In particular, we have shown that PISA can image sub-mm-thick gel at the single-molecule level, achieving attomolar sensitivity. This remarkable sensitivity opens up the possibility for the analysis of single-cell lysates using polyacrylamide gel electrophoresis (PAGE), as even trace amounts of protein from individual cells can be detected.

Here, we present single-cell PAGE-PISA, a highly sensitive strategy for single-cell proteome profiling that integrates PAGE with 3D single-molecule fluorescence imaging using PISA. In this strategy, individual cells are isolated, and proteins are fluorescently labelled, separated by PAGE, and quantified at the single-molecule level. For labelling, we employed *N*-hydroxysuccinimide (NHS)-ester dyes that react with the primary amines of proteins. As the labelling does not rely on antibody, it enables unbiased profiling analysis of the overall cellular proteome without predefined targets, although it does not provide direct identification of the proteins represented in each band. This generates highly sensitive and quantitative electrophoretic band patterns that reflect global protein abundances in individual cells, enabling downstream analyses such as clustering, trajectory inference, and identification of functional cell states solely based on proteome-level variation. To demonstrate the potential of our development, we applied single-cell PAGE-PISA on standard cell lines, PC-3 and U2OS, and explored the temporal changes of cellular proteomes during cardiomyocyte differentiation from human induced pluripotent stem cells (hiPSCs) by tracking their progression through different developmental stages. The resulting proteome profile revealed gradual proteomic changes across single cells that aligned with the progression from pluripotency to differentiated states. Unsupervised clustering and pseudotime analysis further uncovered intermediate subpopulation along the differentiation trajectory, which are often difficult to resolve using transcriptomic analysis or marker-based approaches.

## Results

### Establishment of the single-cell PAGE-PISA workflow

To provide a streamlined proteomic strategy for single cells, we established a complete workflow of single-cell PAGE-PISA: (*1*) cell preparation, (*2*) manual isolation of single target cells, (*3*) one-pot sample preparation in PCR tubes, including cell lysis and protein labelling with dye, (*4*) SDS-PAGE for protein separation, and (*5*) volumetric single-molecule imaging of dye-labelled proteins in polyacrylamide gel using PISA (Fig. 1).

**Figure 1.**
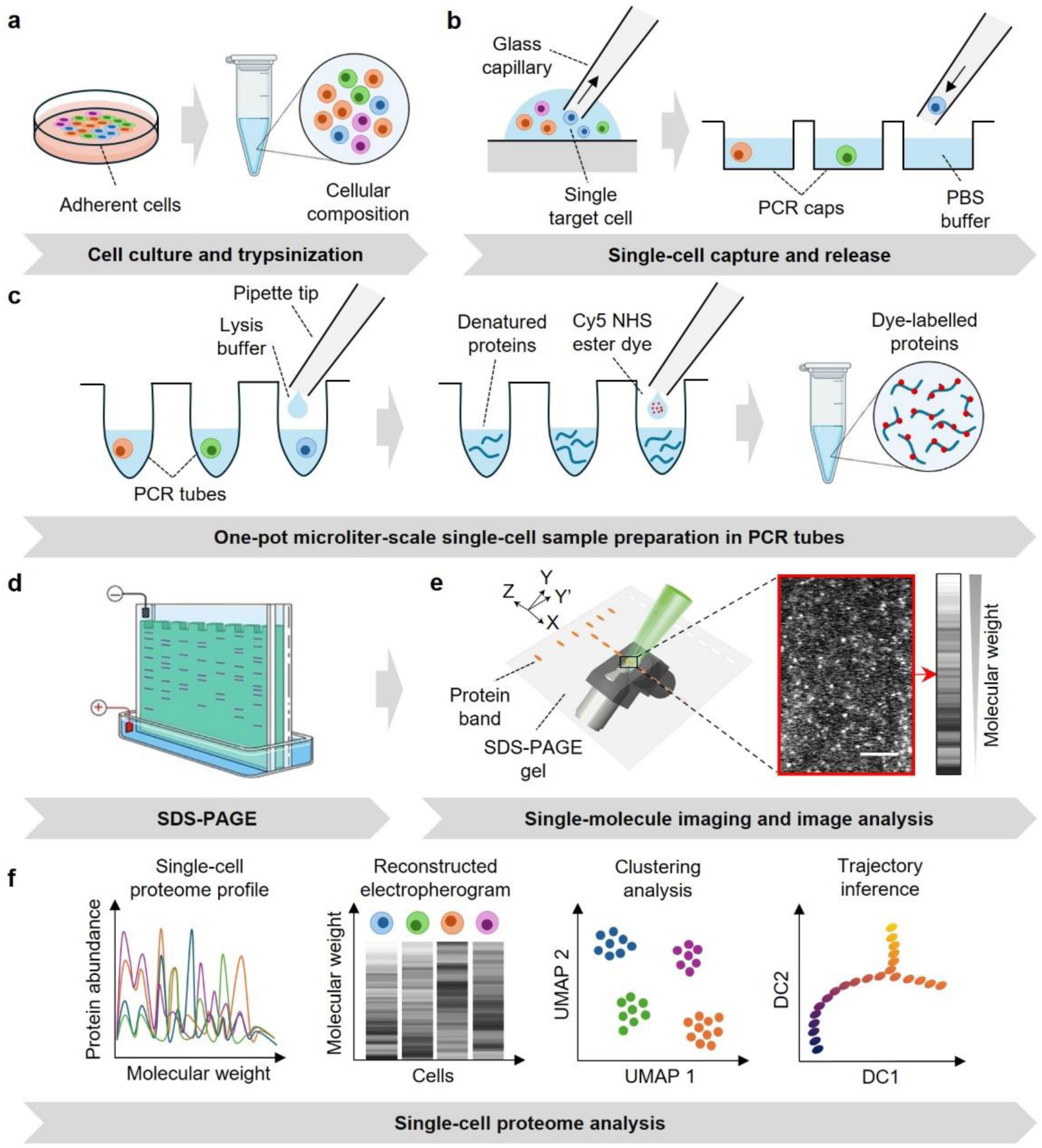
Workflow of single-cell PAGE-PISA. **a** Firstly, cells are dissociated, washed three times, and suspended in PBS at a concentration appropriate for single-cell collection. **b** Then, the single cells are manually isolated from a droplet of cell suspension using an inverted microscope equipped with a cell picker (TOPick 1-cell handling system) and transferred into PCR caps containing PBS. The isolated single cells are spun down to ensure they settle at the bottom of the PCR tubes. **c** After cell lysis, the proteins are labelled with Cy5-NHS ester dye solution at 15°C for 45 minutes. To inhibit the reactivity of excess dye, quencher solution is added to the protein sample, followed by a second incubation at 15°C for 45 minutes. All steps are performed on ice. The molar ratio of the dye to quencher solution used in the single-cell sample preparation is 1:10. **d** The dye-labelled proteins from each single cell are loaded into the well and separated by molecular size using SDS-PAGE. **e** After protein separation, the dye-labelled proteins in the polyacrylamide gel are visualized using PISA with a 647 nm excitation wavelength. Each detected single molecule observed in the PISA image represents a single dye-labelled protein. Fluctuations in the number of single molecules are observed along the migration path, which correlate with protein bands of different abundances. Subsequent image analysis, including background subtraction, denoising filter, and spot detection, enables the reconstruction of a detailed proteome profile for each single cell. Scale bar: 50 µm **f** Schematic diagrams depicting different single-cell proteome analyses that can be achieved with single-cell PAGE-PISA. In (**a**, **c**), the illustration of Eppendorf tube was created with BioRender.com.

In the workflow, cells are first dissociated from the culture dish, washed several times, and suspended in phosphate-buffered saline (PBS) solution (Fig. 1a). Then, a droplet of cell suspension is deposited onto a slide glass, from which the single cells are manually isolated using an inverted microscope equipped with a TOPick 1-cell handling system (i.e., low-binding coated micro-glass needle with a tip diameter of 30 µm and aspirating volume of 100 pL) (Fig. 1b, Supplementary Fig. 1a). The isolated single cells are dispensed into PCR caps containing a droplet of PBS and spun down to ensure the droplet settles at the bottom of the tube. With manual single-cell isolation, we can precisely control cell occupancy by eliminating the possibility of droplets containing multiple or no cells. The entire process of identifying, picking up, and releasing each target cell takes less than two minutes.

Furthermore, we perform single-cell sample preparation on ice without direct contact between pipette tips and protein samples (Fig. 1c). Instead, all reagents, including lysis buffer, protease inhibitor, Cy5-NHS ester dye, and quencher solution, are dispensed along the tube wall without touching the samples directly and mixed by spinning down. This ‘all-in-one’ sample preparation method enhances protein recovery for quantitative single-cell proteome profiling by considerably minimizing potential loss due to surface adsorption and sample transfer.

Next, to separate proteins by molecular weight, we use a commercial 20-well polyacrylamide gel designed for low sample volume with a capacity of up to 8 µL per loading well (Fig. 1d, Supplementary Fig. 1b). The use of a standard polyacrylamide gel with an 8 cm length, instead of preparing a miniaturized gel like the single-cell Western system^26^, improves the separation resolution of the band profiles by resolving proteins with close molecular weights. Meanwhile, the narrow loading well of approximately 4 mm width further concentrates the protein samples within each band along the migration path. As protein bands from single cells cannot be observed by the standard gel imager, we optimized the electrophoresis conditions for single cells (e.g., time, voltage, and current) using the bulk cell lysates in a preliminary experiment. The resulting band profiles from bulk cell lysates, visualized by the standard gel imager allowed us to determine the exact migration pattern and position of each protein band on the polyacrylamide gel (Supplementary Fig. 2a). After single-cell electrophoresis, the gel is carefully cut, transferred onto a film substrate, and placed on a microscopic stage (Supplementary Fig. 1c, d).

Finally, we perform 3D single-molecule imaging using PISA to detect and quantify the dye-labelled proteins, which can be observed as clear diffraction-limited spots during imaging (Fig. 1e). We focus on the middle molecular weight region, approximately from 20‒40 to 100‒150 kDa, which corresponds to a 5 cm gel length (Supplementary Fig. 1d). The region of interest is determined by the size of the metal stage and the balance between measurement time and sufficient proteome information. The single-molecule imaging is conducted with a scan speed of 80 µm/s, resulting in approximately 10 minutes of imaging time per single cell. Fluctuations in the number of dye-labelled proteins are observed along the migration path during imaging, which reflects protein bands with differing protein abundances across molecular weights (Supplementary Fig. 2c, Supplementary Movie 1). Detecting and quantifying the dye-labelled proteins at the single-cell level allows us to gain insights on protein expression profiles, cell type classification, and pseudotime analysis (Fig. 1f).

To evaluate the sensitivity and quantitative accuracy of the PISA measurement, we imaged and quantified the number of dye-labelled proteins (referred to as protein count) from highly diluted bulk HeLa and U2OS cell lysates, equivalent to a single-cell concentration. Note that the bulk cell lysates were diluted to single-cell level based on the estimated initial number of the cell suspension. We found that the reconstructed electropherogram obtained by single-molecule imaging was in good agreement with the SDS-PAGE band profile visualized by the standard gel imager (Supplementary Fig. 2b and Supplementary Fig. 2a, respectively). Quantitative analysis revealed that the protein counts across molecular weights were consistent in three technical replicates, with a median coefficient of variation (CV) of 6.5% and 7.8% for HeLa and U2OS, respectively (Supplementary Fig. 2d, e). The analytical reproducibility was assessed by computing the Pearson correlation of the protein counts across molecular weights between replicates, which yielded coefficients larger than 0.96 (Supplementary Fig. 2f). To assess the linearity of the method, we performed an experiment using highly diluted bulk cell lysates equivalent to 1, 2, 4, and 8 cells. Bulk cell lysates were used to minimize the variability from cell-to-cell differences in protein content or cell cycle stages, thus ensuring controlled and consistent protein input. Our result demonstrated excellent linearity (R² = 0.998) between cell equivalents and total protein counts (Supplementary Fig. 2g, h). The ability to maintain linearity down to single-cell equivalent shows robust quantitative accuracy and great detection sensitivity of our system, even in highly diluted samples.

Lastly, to determine the sensitivity of single-cell PAGE-PISA to detect low-abundance proteins, we conducted a lysate spike-in experiment using transferrin (77 kDa) as a model protein and evaluated its detectability within a complex cell lysate environment (Supplementary Figure 3a). After subtraction of the negative control, we successfully detected as little as 5 fg of transferrin, corresponding to approximately 7 × 10⁴ protein molecules (Supplementary Figure 3b). These results demonstrate that single-cell PAGE-PISA can reliably detect and quantify proteins present at copy numbers ranging from 10⁴–10⁵ within a complex cell lysate background.

### Single-cell proteome profiling of mammalian cells

Next, we performed single-cell proteomic analysis using single-cell PAGE-PISA on two different tumour cell lines, U2OS and PC-3 (Fig. 2). The single cells were manually isolated, prepared in PCR tubes, and the dye-labelled proteins were separated by SDS-PAGE and imaged by PISA. The imaging was performed at the middle molecular weight region, covering approximately 40‒50% of the cellular proteome. The electropherogram was reconstructed from the PISA images to provide a comprehensive view of the protein expression profiles across multiple single cells (10 biological replicates for each cell type) (Fig. 2a). In practice, our single-cell PAGE-PISA could quantify dye-labelled proteins from a 2 mm^3^ gel volume (0.2 × 0.2 × 50.0 mm^3^). This translates to 10^3^‒10^4^ dye-labelled proteins per band, corresponding to approximately 2 fg, and a total of 10^5^ dye-labelled proteins per single cell. Ideally, imaging the entire migration path (4.0 × 1.0 × 50.0 mm^3^) could theoretically increase these numbers to 10^5^‒10^6^ dye-labelled proteins per band, which corresponds to approximately 200 fg, and a total of 10^7^ dye-labelled proteins in a single mammalian cell (Fig. 2b, c, Supplementary Fig. 4a). This detection sensitivity far surpasses that of conventional staining methods for SDS-PAGE like Coomassie Brilliant Blue (CBB) and silver staining, with detection thresholds of 8‒10 ng^30^ and 0.1‒0.5 ng^31,32,33^ per protein band, respectively. Thus, these methods were insufficient to visualize protein bands from single-cell samples, unlike single-cell PAGE-PISA which can detect protein bands with atto-gram amounts^29^. Thereby, the detection sensitivity was improved by 3‒5 orders of magnitude over silver staining and CBB, enabling the detection of even 1% of the total protein abundance.

**Figure 2.**
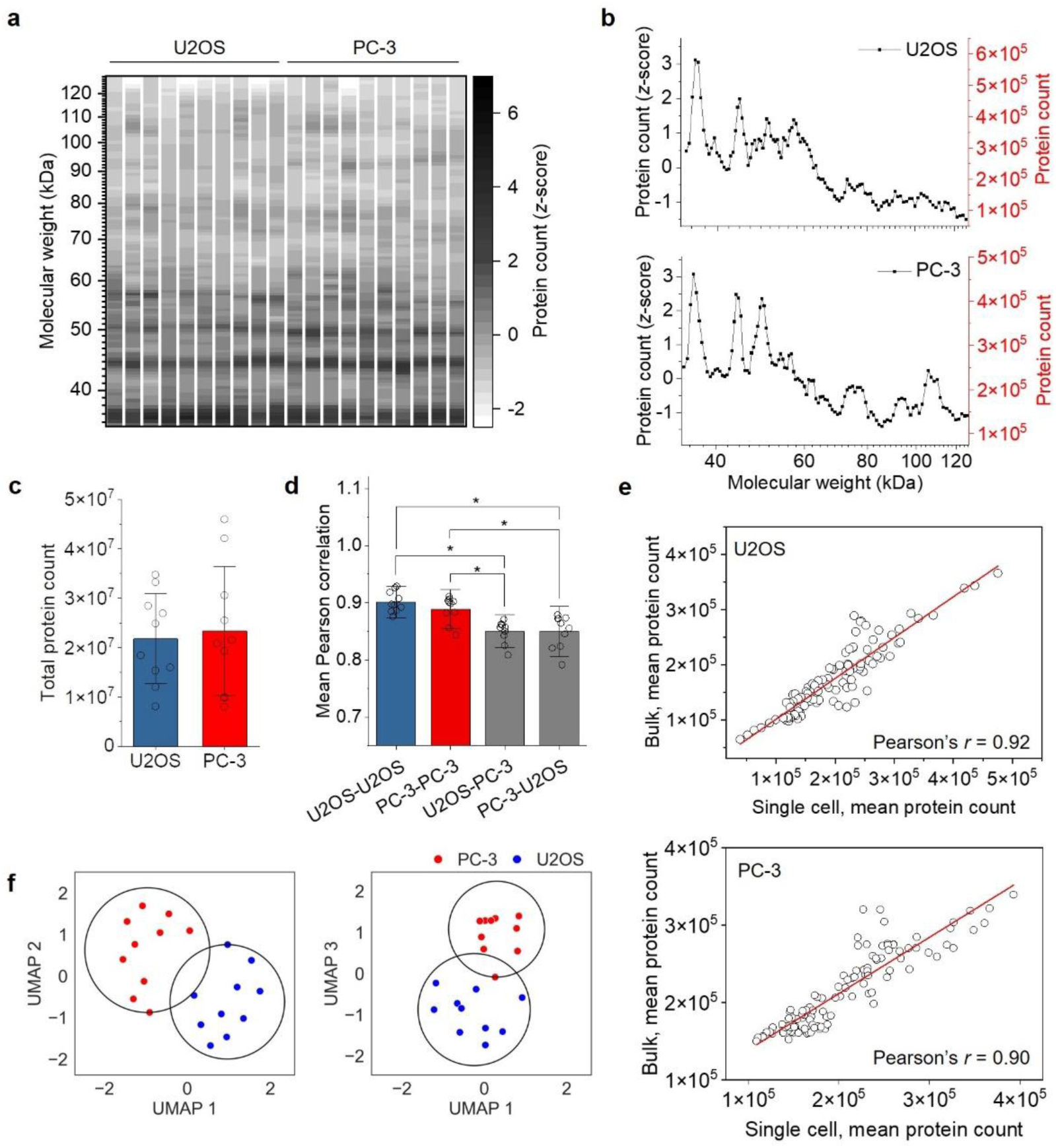
Single-cell PAGE-PISA of single U2OS and PC-3 cells. **a** Reconstructed single-cell electropherogram of U2OS and PC-3 cells obtained by single-cell PAGE-PISA. The electropherogram was reconstructed based on the standardized protein counts from 35‒128 kDa. **b** Representative proteome profile of single U2OS and PC-3 cells. The *y*-axes represent the standardized (left) and raw (right) protein counts from a single cell across molecular weights, respectively. **c** Total protein count for each cell type. **d** Mean Pearson correlation of protein counts across molecular weights between single cells, within and across cell types. Error bars represent ± the standard deviation of the mean. **e** Comparisons of the mean protein count between single cells and technical replicates of highly diluted bulk cell lysates equivalent to single-cell level. The red lines represent the line of best fit, showing a positive correlation between single-cell and bulk measurement. **f** Two-dimensional UMAP projections based on electrophoretic protein band profiles from individual U2OS (blue) and PC-3 (red) cells, shown as UMAP 1 vs. UMAP 2 and UMAP 1 vs. UMAP 3. Clusters (grey region) were identified using *k*-means. Number of biological replicates (single cells): *n* = 10 for U2OS and *n* = 10 for PC-3, number of technical replicates (bulk cell lysates): *n* = 3 for U2OS and *n* = 3 for PC-3 cells. Statistical significance was determined by the one-way Welch’s ANOVA followed by the Games-Howell post-hoc test. \**p* <= 0.05

Compared to the estimated 6 × 10^9^ protein copies per HeLa cell^34^, the total protein copies quantified by single-cell PAGE-PISA fall short by two orders. The difference between practical and theoretical values may stem from technical limitations during quantification. For instance, residual dyes that are not completely quenched may weakly interact with the polyacrylamide gel during electrophoresis, causing a slight increase in the overall background signal (Supplementary Fig. 5a, b, Supplementary Movie 2). As a result, some dye-labelled protein molecules that are not distinctive from the background may fall below the detection threshold and failed to be quantified (Supplementary Fig. 5c). Furthermore, if two dye-labelled protein molecules are within the optical diffraction limit (i.e., a few hundred nanometers), especially in highly dense protein bands, they cannot be resolved and will be detected as a single molecule due to the diffraction limit. Nevertheless, the strong mean Pearson correlation of protein counts across molecular weights between single cells from the same cell types (*r* > 0.89) demonstrates great reproducibility and a prominent level of quantitative accuracy of single-cell PAGE-PISA (Fig. 2d, Supplementary Fig. 4b).

Next, we conducted a detailed peak fitting analysis to determine the peak capacity and mass resolution of single-cell PAGE-PISA. We identified 21 distinct Gaussian peaks between 35 and 128 kDa in a representative single-cell proteome profile (Supplementary Fig. 6a). For proteins in the 35–60 kDa range (peaks #1–8), the average mass resolution was 1.34 kDa (FWHM) (Supplementary Fig. 6b). For proteins in the 60–128 kDa range (peaks #9–21), the resolution declined to an average of 2.69 kDa, likely due to reduced gel migration efficiency for large proteins.

We further assessed the accuracy and reliability of our single-cell measurement by comparing the single-cell protein expression profile to those of highly diluted bulk cell lysates equivalent to a single-cell concentration from the same cell types. We averaged the protein counts across multiple single cells (*n* = 10) and technical replicates of highly diluted bulk cell lysates (*n* = 3), then compared their mean protein count across molecular weights (Supplementary Fig. 4c). We observed a strong Pearson correlation of mean protein counts between single cells and highly diluted bulk cell lysates for both U2OS and PC-3 cells (*r* > 0.90) (Fig. 2e). The observed correlation showed that the single-cell protein quantification by single-cell PAGE-PISA was in good agreement with that from bulk cell lysates diluted to the single-cell level, indicating the robustness and reliability of our single-cell measurement.

Lastly, we visualized the 2D projections of single U2OS and PC-3 cells, specifically in UMAP 1 vs UMAP 2 and UMAP 1 vs UMAP 3 plots (Fig. 2f). Our results showed clear separation between the two cell lines, indicating that the clustering captured meaningful biological differences and was not confined to higher-order dimensions. This demonstrates that single-cell PAGE-PISA can effectively identify and classify different cell types solely based on their proteome.

### Single-cell proteome profiling of cardiomyocyte differentiation from human induced pluripotent stem cells

The ability of hiPSCs to self-renew and differentiate into various cell types and tissues has garnered significant interest, especially in the pursuit of heart regenerative therapies^35,36^. Numerous studies have shown that protein expressions of hiPSC-derived cardiomyocytes (hiPSC-CMs) exhibit dynamic fluctuations and temporal changes during differentiation and maturation^37,38,39^.

To highlight the potential applicability and versatility of single-cell PAGE-PISA in detecting global proteome changes during dynamic processes, we performed single-cell proteome profiling analysis on hiPSCs as they progressed from pluripotency through stage-specific transitions during cardiomyocyte differentiation (Fig. 3a). We observed cardiac contraction in hiPSC-CMs as early as day 7, indicating the successful differentiation process. To improve cardiomyocyte purity in culture, the hiPSC-CMs were carefully dissociated and transferred to a freshly coated plate between day 10 and 12. Cells were harvested on different days, including day 0 (hiPSCs), 16, and 30 (hiPSC-CMs), referred to as D0, D16, and D30, respectively. In total, 49 single cells were collected, prepared, and analyzed with single-cell PAGE-PISA (*n* = 12, 14, and 23 for D0, D16, and D30, respectively) (Fig. 3b).

**Figure 3.**
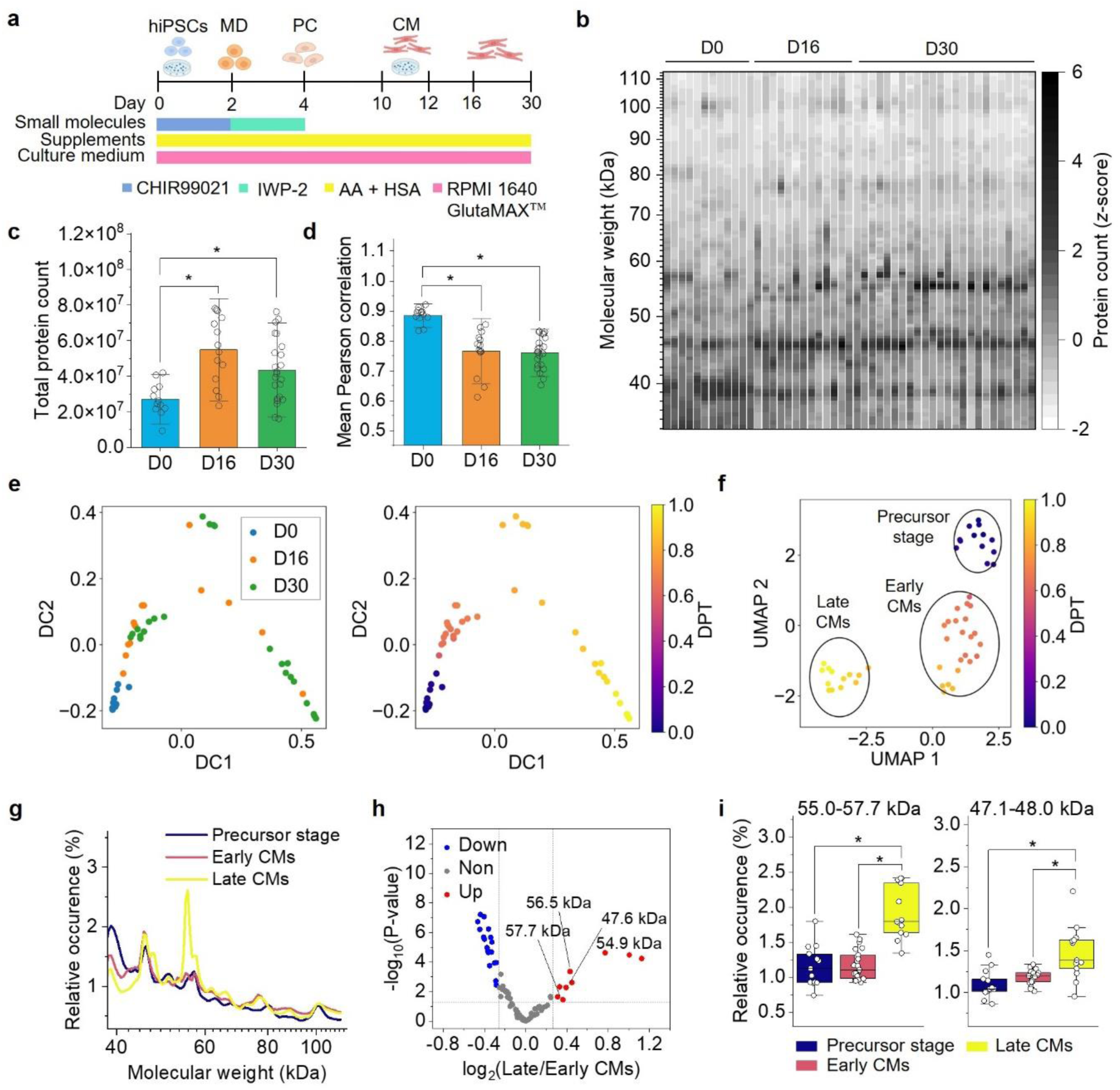
Application of single-cell PAGE-PISA to cardiomyocyte differentiation from human induced pluripotent stem cells. **a** Schematic protocol of cardiomyocytes transitioning from pluripotent state towards cardiac lineage. hiPSCs: human induced pluripotent stem cells; MD: mesoderm; PC: progenitor cells; CM: cardiomyocytes; AA: ascorbic acid; HSA: human serum albumin. The culture dish represents the day when hiPSC-CMs were transferred to a freshly coated plate. Illustrations were created with BioRender.com. **b** Single-cell electropherogram of hiPSCs (D0) and hiPSC-CMs (D16 and D30) were reconstructed based on the standardized protein counts from 35‒112 kDa. **c** Total protein count for each sampling day. **d** Mean Pearson correlation of protein counts across molecular weights between single cells, within and across sampling days. Error bars represent ± the standard deviation of the mean. Number of biological replicates: *n* = 12 for D0, *n* = 14 for D16, and *n* = 23 for D30. **e** Diffusion maps showing the developmental trajectory from pluripotency to differentiated states. The single cells were coloured according to their sampling day (left) and DPT (right). The trajectory starts from the root cell (blue) that is located at the left terminal and progresses towards the most differentiated cell (yellow). DC and DPT refer to diffusion components and diffusion pseudotime, respectively. **f** UMAP analysis with *k*-means clustering of the DPT-assigned single cells. Three clusters were identified, representing different developmental stages, precursor stage (*n* = 13), early cardiomyocytes (*n* = 23) and late cardiomyocytes (*n* = 13). **g** Average single-cell proteome profile for each developmental stage. **h** Volcano plot showing upregulated and downregulated protein bands in late cardiomyocytes compared to early cardiomyocytes. Differential protein bands were determined by Welch’s two-sided, two-sample *t*-test (*p* < 0.05, fold change > 1.2). **i** Comparisons of the quantitative expression levels of two protein bands, 55.0‒57.7 kDa and 47.1‒48.0, across three developmental stages. The box plots show median values (central line), interquartile range (box edges), and whiskers extending up to 1.5 times the interquartile range. Statistical significance was determined by the one-way Welch’s ANOVA followed by the Games-Howell post-hoc test. Individual data points are overlaid. \**p* <= 0.05

Quantitative analysis of single hiPSC-CMs from D16 and D30 revealed 1.5‒2.0 times higher total protein abundance than D0, likely due to the increased cell size and complexity in specialized cells (Fig. 3c). In addition, we observed a strong mean Pearson correlation of protein counts across molecular weights among D0 single cells (*r* = 0.88), implying a consistent single-cell protein expression levels in the undifferentiated state (Fig. 3d). Meanwhile, the mean correlation of protein counts decreased by approximately 13.6% in both D16 and D30 single cells, suggesting stochastic occurrences in individual cells during cardiomyocyte differentiation. Using all the standardized protein counts from 35‒112 kDa, principal component analysis (PCA) separated the cells in a time-wise manner along PC1, which accounts for 44.1% of the total variance (Supplementary Fig. 7a). Notably, the data points from different batches did not form distinct clusters in the PCA plot, indicating that batch effects were negligible and did not confound the temporal separation.

We next sought to investigate the cell progression from pluripotency towards differentiated states by pseudotemporal ordering of the dynamic cells across a developmental lineage^40^. Briefly, cells were presented on a diffusion map based on the cell-to-cell transition probabilities, positioning those with higher transition probabilities closer to each other (Fig. 3e, left). The trajectory was inferred by computing diffusion pseudotime (DPT) for each cell relative to the root cell, often considered the starting point of the differentiation process, and cells were ordered along the pseudotemporal trajectory to track their developmental progress during cardiomyocyte differentiation (Fig. 3e, right). As anticipated, the D0 single cells were in close proximity to one another on the diffusion map, indicating homogeneous protein expression profiles among the single cells. In contrast, cells from D16 and D30 appeared more spread out across the developmental trajectory, with D16 and D30 primarily dominating the early and terminal end of the trajectory, respectively. This reflects the temporal dynamics of proteome changes as cells differentiate from early to later stages. We performed UMAP analysis on DPT-assigned single cells and identified three clusters (Fig. 3f, Supplementary Fig. 7b). Precursor stage (*n* = 13) consisted predominantly of D0 cells (*n* = 12, 92.3%), while early (*n* = 23) and late cardiomyocytes (*n* = 13) were mainly composed of D16 and D30 cells (*n* = 10 and *n* = 10, 43.5% and 76.9%, respectively).

We further visualized the single-cell proteome profiles across different developmental stages to identify differentially regulated protein bands (Fig. 3g, Supplementary Fig. 8a). By comparing the protein expression differences between early and late cardiomyocytes, we identified two protein bands that were differentially regulated in late cardiomyocytes (Fig. 3h, Welch’s two-sided, two-sample *t*-test, *p* < 0.05, fold change > 1.2). Notably, the protein bands at 55.0‒57.7 kDa and 47.1‒48.0 kDa were significantly expressed in late cardiomyocytes compared to the precursor stage and early cardiomyocytes (Fig. 3i). However, due to the lack of reported cardiac-specific proteins within these molecular weight ranges, we hypothesize that the observed expression may be associated with certain biological processes which are crucial for cardiomyocyte development. To support our notion, we compared our single-cell proteome data to the single-cell transcriptome data obtained from a public database^41^. We averaged the transcript counts across single cells for each sampling day: D0 (*n* = 9,146), D15 (*n* = 2,897), and D30 (*n* = 3,294), visualized their mean transcriptome profiles, and identified upregulated genes in D30 (Supplementary Fig. 8b, c, Welch’s two-sided, two-sample *t*-test, *p* < 0.05, fold change > 1.4). Among the differentially upregulated genes with the highest transcript count within 55.0‒57.7 kDa and 47.1‒48.0 kDa were *ENO1* (47.2 kDa), *ATP5B* (56.5 kDa), and *MT-CO1* (57.0 kDa) (Supplementary Fig. 8d). In addition, a previous study has shown that both MT-CO1 and ATP5B proteins were significantly expressed in the cavities of cardiac tissue^42^, consistent with their role in energy production during cardiac development. Considering these genes are quantitatively abundant and significantly expressed at both transcriptome and proteome levels, we believe that their expression levels are reflected in single-cell PAGE-PISA.

### Comparative analysis between single-cell proteome and transcriptome during cardiomyocyte differentiation

As proteome and transcriptome measurements are not always correlated^5,6,7,8,9^, we next compared the extent to which our single-cell proteome data obtained by single-cell PAGE-PISA are in agreement with reported single-cell RNA sequencing (scRNA-seq)^41^ during cardiomyocyte differentiation. The scRNA-seq data was chosen as the sampling days when the single cells were collected closely matched our single-cell PAGE-PISA. We analyzed both datasets with an equal sample size (*n* = 49) for accurate comparison. Three random samples (RS) were generated from the scRNA-seq data from 37‒126 kDa, with each RS consisting of 49 randomly selected single cells from three different sampling days (*n* = 12, 14, and 23 for D0, D15, and D30, respectively). UMAP analysis revealed a consistent spatial distribution of the cells at both proteome and transcriptome levels (Fig. 4a). The D0_protein_ and D0_RNA_ populations formed a distinct cluster, with cells in close proximity to each other, implying that both populations exhibit stable developmental states with consistent expression profiles. Meanwhile, D16_protein_, D30_protein_, D15_RNA_, and D30_RNA_ populations displayed two separate clusters that were distinctive from D0 but did not otherwise cluster with respect to their sampling days. These results indicate that although D16_protein_, D30_protein_, D15_RNA_, and D30_RNA_ populations share similar expression profiles on UMAP space, they reflect transitional states or different developmental stages during cardiomyocyte differentiation.

**Figure 4.**
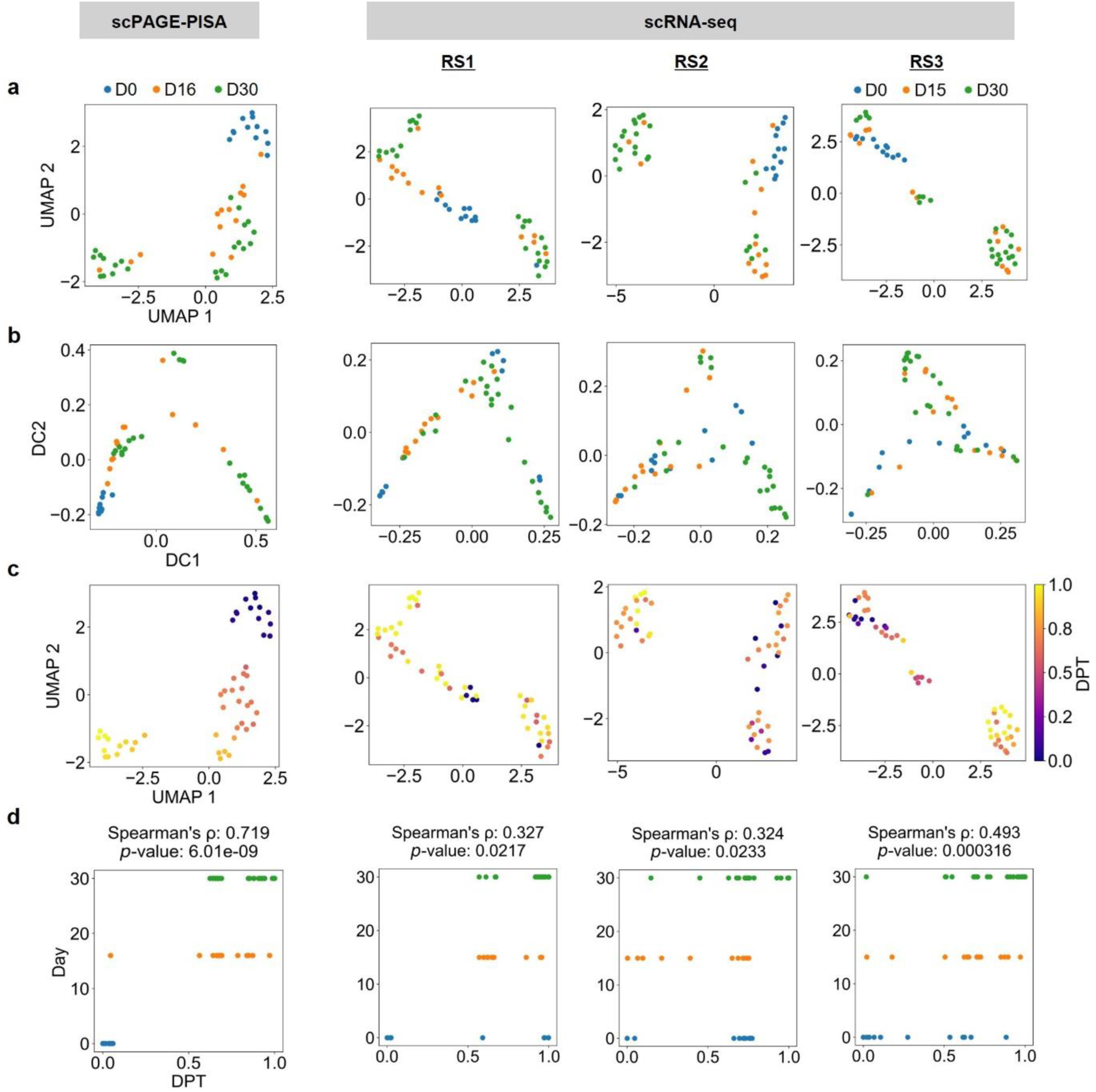
Comparative analysis between single-cell proteome and transcriptome during cardiomyocyte differentiation. **a** UMAP projection of single cells based on the overall similarity of gene or protein expression. **b** Diffusion maps showing single cells arranged along a pseudotemporal trajectory based on the transition probabilities between two cells, positioning those with higher transition probabilities closer to each other. **c** UMAP projection of single cells after DPT assignment. **d** Correlation between sampling day and DPT. For scRNA-seq data, three RS were generated by randomly selecting 49 single cells from the entire dataset. In total 10,484 genes were used for the analysis, corresponding to proteins with molecular weights within 37‒126 kDa. DC refers to diffusion components, DPT refers to diffusion pseudotime, and RS refers to random sample. Number of biological replicates for single-cell PAGE-PISA: *n* = 12, 14, and 23 for D0, D16, and D30, respectively. Number of biological replicates for scRNA-seq: *n* = 12, 14, and 23 for D0, D15, and D30, respectively.

We expect the spatial distribution of these states can be better resolved on a pseudotemporal scale. Hence, we ordered the dynamic cells along a developmental trajectory from pluripotency to differentiated states at both proteome and transcriptome levels (Fig. 4b, Supplementary Fig. 9b, c). As expected, the majority of D30_protein_ and D30_RNA_ single cells were located at the terminal end of the trajectory, distal from the root cell, signifying their progression into a fully differentiated state. While D0_protein_ population clustered closer to the root cell on the diffusion map, we observed greater variations of D0_RNA_ population in all three RS, with cells sparsely distributed across the developmental trajectory. The D0_RNA_ population, however, can be clearly distinguished from D15_RNA_ and D30_RNA_ populations when increasing the sample size from 49 to 3,000 single cells (Supplementary Fig. 9a). This emphasizes the need for a larger sample size of scRNA-seq data^43^ to capture the full spectrum of biological variability.

Lastly, we sought to investigate how well the inferred DPT correlates with the sampling day (referred to as real-time) during cardiomyocyte differentiation at both proteome and transcriptome levels. To this end, we visualized the DPT-assigned single cells on UMAP and assessed the correlation between the DPT and real-time (Fig. 4c, d). We observed a strong concordance between DPT_protein_ and real-time_protein_ (Spearman’s rho = 0.72), while relatively lower concordance between DPT_RNA_ and real-time_RNA_ in all three RS (Spearman’s rho < 0.49) (Fig. 4d). These results demonstrate that the directional changes of the developmental states at proteome level are generally consistent whether determined by DPT or real-time and that both recapitulate the expected temporal changes during cardiomyocyte differentiation. While several D15_RNA_ and D30_RNA_ cells displayed a wide dynamic range of DPT values due to the different progression rate during the differentiation process, it is more pronounced in D0_RNA_ populations, suggesting the presence of subpopulations at the pluripotent state, consistent with previous reports^44,45^. The observed disparities could be attributed to the difference in stabilities and half-lives of RNA and protein. By comparison, RNA has a relatively shorter half-life and is more susceptible to degradation than proteins^46^. In turn, it directly impacts the RNA abundance and leads to temporal variations during sampling. In contrast, long protein half-lives and greater stability have led to more consistent expression levels, which can accurately reflect the cellular states and functions.

Overall, these findings demonstrate the ability of single-cell PAGE-PISA to provide a direct and robust depiction of developmental status during dynamic processes, even with minimal sample sizes. Its ability to resolve developmental trajectories from as few as 49 cells not only is particularly promising for studies of rare cell populations or samples with limited availability, but it also opens up new avenues in various research fields, including developmental biology, regenerative medicine, and disease modeling. Together, single-cell PAGE-PISA serves as a powerful complementary approach to scRNA-seq by bridging the gaps between transcriptome and proteome and providing a more comprehensive understanding of cellular dynamics at the single-cell level.

## Discussion

In this present work, we introduced single-cell PAGE-PISA capable of highly sensitive and quantitative profiling of a wide range of proteins in individual cells. This system was realized through the integration of PAGE for protein separation based on molecular sizes with 3D single-molecule imaging using PISA for protein quantification. The workflow is streamlined and benchtop-compatible, using microliter-scale sample volumes in standard PCR tubes that are easily handled with pipettes, eliminating the need for specialized robotic systems.

With single-cell PAGE-PISA, we detected 21 protein bands per single cell and quantified 10^7^ molecules covering approximately 1% of the total proteome per cell. This proportion can be further improved to over 10% by using brighter dyes such as SeTau-647^47^ or advanced spot-recognition algorithms, including deep learning^48^ (Supplementary Fig. 10). With further improvements in separation resolution and labelling efficiency, we anticipate achieving detection sensitivity down to 10^2^–10^3^ molecules per species, enabling the analysis of low-abundance proteins.

A key advantage of single-cell PAGE-PISA is its ability to uncover functional subpopulations that may not be distinguishable using conventional markers or transcriptomic profiling. For example, heterogeneous T cell populations often exhibit diverse activation states that are not easily resolved by known surface markers used in cell sorting^49,50^. By capturing the full proteomic content of individual cells, our method enables the identification of rare or intermediate states based on functional protein features. Another strength is its ability to resolve modification-specific and structural proteoforms that are often inaccessible to MS-based approaches due to sample loss, limited enrichment efficiency, or disruption of protein complexes during sample preparation. This is enabled by integrating Phos-tag^51^ PAGE or native PAGE, which permits direct detection of phosphorylated isoforms and native protein complexes at the single-cell level—information critical for understanding cellular decisions governed by signaling assemblies such as TNF receptor complexes^52^. Furthermore, by labelling specific proteins of interest in situ, this platform could be extended to spatially resolved subcellular proteomics, enabling the analysis of organelle-associated proteins while preserving intracellular localization.

Although chip-based electrophoresis, as implemented in single-cell Western blotting^26^, and capillary electrophoresis (CE) can, in principle, be adapted for proteome analysis using PISA, both approaches face significant limitations for single-cell proteome profiling. The format used in single-cell Western offers high-throughput analysis of over 1,000 cells but suffers from limited separation resolution due to its short migration distance (∼1 mm), resulting in overlapping bands and reduced quantification accuracy. CE systems, such as the PA 800 Plus (Sciex), can provide higher-resolution separation due to longer capillary lengths. However, their workflows are inherently sequential and typically require 1–2 hours per cell, which imposes a major constraint on throughput. Taken together, single-cell PAGE-PISA offers a practical and effective balance of resolution, sensitivity, and throughput for comprehensive proteome profiling at the single-cell level.

Although single-cell PAGE-PISA currently requires approximately 7 hours to analyze 20 cells simultaneously, scaling up to thousands of cells will likely be necessary to address broader biological questions. We anticipate that such expansion will be feasible through the use of automated single-cell collection (e.g., fluorescence-activated cell sorting or automated live imaging and picking systems such as ALPS^53^), high-throughput robotic reagent handling, and rapid, parallel electrophoresis (e.g., 10-minute runs at 400 V using bullet PAGE from Nacalai Tesque). Although imaging currently limits the overall throughput, our preliminary tests demonstrate that scan speeds can be increased up to 400 µm/s without loss of accuracy^29^. The integration of high-speed CMOS cameras with larger fields of view could further accelerate data acquisition. Overall, scaling up to thousands of cells is estimated to require approximately 30 hours from cell isolation to PISA imaging. Since single-cell PAGE-PISA uses only commercially available materials, the cost per cell is estimated at $0.60–1.00 and can be reduced to ∼$0.11 using homemade gels, making it a highly cost-effective alternative to other single-cell proteomics methods.

Currently, a major limitation of single-cell PAGE-PISA is the inability to assign molecular identities to individual protein bands, which limits the biological interpretability of the protein expression profiles and constrains the applicability for precise cell phenotyping. To address this limitation, one potential strategy is to incorporate multiplexed protein identification directly within the gel using antibodies or multicolor tags, similar to In-Gel Western^54^. Additionally, combining single-cell PAGE-PISA with fluorescent antibody-based sorting via FACS and conventional MS can help associate single-cell proteome profiles with specific protein identities. This integrative approach enables molecular annotation of phenotypically meaningful bands and enhances the ability to interpret cell state differences observed in single-cell PAGE-PISA.

In conclusion, we have established single-cell PAGE-PISA and demonstrated its potential for ultra-sensitive single-cell proteome analysis on diverse cell types, from steady-state to differentiated cells. We strongly believe that our system can help researchers narrow down the number of potential protein species in each band at single-cell level prior to immunofluorescent imaging or MS. By focusing on a smaller subset of proteins, this approach not only will help to reduce the time and resources required for downstream analysis but also minimize the need for extensive antibody libraries, making target protein identification more cost-effective for future biomarker discovery.

## Materials and methods

### Cell culture

U2OS, HeLa (RCB0007), and PC-3 (RCB2145) cells were obtained from the RIKEN cell bank. The U2OS and HeLa cells were maintained in Dulbecco’s modified eagle medium (DMEM) (Thermo Fisher Scientific, 10566-016) supplemented with 10% fetal bovine serum (FBS) (Corning, 35-079-CV). The PC-3 cells were maintained in RPMI 1640 medium (Thermo Fisher Scientific, 11875-093) supplemented with 10% FBS. Human induced pluripotent stem cells (hiPSCs) were purchased from RIKEN BRC (HPS4290:201B7-Ff) and maintained in mTeSR^TM^ Plus medium (STEMCELL Technologies, 100-0276). All cell lines were maintained in a 5% CO_2_ incubator at 37°C.

### Cardiomyocyte differentiation protocol

Cardiomyocyte-directed differentiation from hiPSCs was performed and modified based on the previous protocol^55^. Four days before cardiomyocyte differentiation, the hiPSCs were dissociated using TrypLE^TM^ Select CTS^TM^ (Gibco, A12859-01) and cultured in a matrigel-coated 24-well plate at a density of 5 × 10^4^ cells/well in mTeSR^TM^ Plus. On day 0, the hiPSCs were washed with phosphate-buffered saline (PBS) (Nacalai-tesque, 14249-95) and treated with differentiation medium (RPMI 1640, GlutaMAX™ (Gibco, 61870036) containing 500 µg/mL human serum albumin (HSA) (Wako, 010-27601) and 213 µg/mL L-ascorbic acid 2-phosphate (Nacalai-tesque, 13571-56)) supplemented with 6 μM CHIR99021 (Wako, 252917-06-9). On day 2, the medium was replaced with fresh differentiation medium supplemented with 5 μM IWP-2 (Wako, 686770-61-6). On day 4, the medium was replaced with a fresh differentiation medium without supplemental inhibitors. Medium exchange was performed every two days. Beating was first observed on day 7.

### Bulk cell lysate sample preparation

Cells were harvested, washed with PBS three times, and counted using an automatic cell counter (TC20, BioRad) to obtain a concentration of 1.5 × 10^3^ cells/µL. Approximately 50 µL of lysis buffer (50 mM borate (Nacalai-tesque), 1% Tween 20 (Sigma-Aldrich, 9005-64-5), 1% sodium dodecyl sulfate (SDS) (Wako, 192-13981), adjusted at pH 8.0) and 5 µL of protease inhibitor (Nacalai-tesque, 25955) were added to 50 µL of cell suspension. After cell lysis, proteins were labelled with Cy5-NHS ester dye (AAT Bioquest, 151) to a final concentration of 100 µM and incubated at 15°C and 2000 rpm for 45 minutes. The dye-labelled proteins were washed with PBS three times in a centrifugal filter unit (10K) (Millipore, UFC5010BK) and centrifuged at 10,000 × g and 4°C for 10 minutes.

Approximately 50 µL of protein sample was collected after the final wash and stored at-20°C.

### Lysate spike-in experiment

Approximately 15,000 cell lysates were spiked with 75 ng, 7.5 ng, and 0.75 ng of purified transferrin (Fujifilm Wako, 205-18084), lysed in the presence of protease inhibitor, and labelled with Cy5-NHS ester dye to a final concentration of 100 µM. The samples were washed with 20 mM borate buffer three times in a centrifugal filter unit (10K) and centrifuged at 10,000 × g and 4°C for 10 minutes. For single-molecule imaging, the lysate spike-in samples were diluted to obtain cell lysates containing 500 fg, 50 fg, and 5 fg of transferrin. Negative control was prepared containing only bulk cell lysates without exogenous protein.

### Single-cell isolation

A droplet of cell suspension was deposited onto a glass slide (Matsunami, S1111) that was placed on the stage of an inverted microscope (CKX41, Olympus) equipped with a TOPick 1-cell handling system, composed of a touch panel, micro liquid pump, and controller (YODAKA Co., Ltd.). The single cells were manually isolated using a 30 µm G-tip low-binding coated micro-glass needle (YODAKA Co., Ltd.) with a minimum handling volume of 100 pL. The isolated single cells were transferred to the PCR cap containing 1 µL of PBS, spun down, and stored at-80°C.

### Single-cell sample preparation

Manually isolated single cells were lysed on ice with 0.6 µL of lysis buffer and 0.4 µL of protease inhibitor. Proteins were labelled with 0.4 µL of Cy5-NHS ester dye to a final concentration of 1 µM and incubated at 15°C and 2000 rpm for 45 minutes. To quench the reactivity of excess unreacted dye, 0.4 µL of Tide Quencher™ 5WS amine (AAT Bioquest, 2076) was added to the protein sample to a final concentration of 10 µM. The quencher was added not only to stop the reactivity of NHS ester but to effectively quench the fluorescence of unreacted Cy5-NHS ester dye. As such, it omits the need for common dye removal methods such as gel filtration or desalting columns, which often result in significant protein loss. For efficient protein labelling, the molar ratio of dye to quencher solution used in this protocol was 1:10. To avoid protein loss, all reagents were dispensed along the walls of the PCR tubes, spun down, and stored at-80°C. Negative control was prepared similar to the single-cell sample but containing only PBS buffer, lysis buffer, protease inhibitor, dye, and quencher solution.

### SDS-PAGE

Electrophoresis was performed using two different polyacrylamide gels depending on the experimental purposes. For standard gel imaging (Fujifilm, LAS 4000), the dye-labelled protein samples from 7.5 × 10^3^ cell lysates were mixed with 4× SDS sample buffer (240 mM Tris-HCl, 40% glycerol stock (Nacalai-tesque, 17045-65), 8% SDS (Nacalai-tesque, 31606-75), and β-mercaptoethanol (Nacalai tesque, 21438-82), adjusted at pH 6.8), heated at 95°C for two minutes, and separated using a 12-well, 5‒20% precast polyacrylamide gel (Bio-craft, #SDG-571). The loading volume was 12 µL. For single-molecule imaging, the dye-labelled protein samples were prepared, heated at 95°C for two minutes, and separated using a 20-well, 5‒20% precast polyacrylamide gel (Bio-craft, #SDG-576). A total volume of 8 µL was loaded for highly diluted bulk cell lysates equivalent to single-cell level and 5 µL for single-cell lysates, both containing 4× SDS sample buffers. To avoid diffusion of free Cy5-NHS ester dye during protein migration, the samples were applied and separated by two loading wells. All gels were irradiated under UV overnight to remove autofluorescence signals that could contribute to the background noise. Electrophoresis was performed at 250 V and 30 mA for 70 minutes.

### Single-molecule imaging

After SDS-PAGE, the polyacrylamide gel was cut vertically into a 5 cm length, covering the majority of the protein bands in the middle molecular weight region, and placed on a UV-irradiated fluorinated ethylene propylene (FEP) film (Daikin Chemical, NF-0025). To avoid gel desiccation and reduce the reflection of excitation illumination on the gel surface during observation, two blank polyacrylamide gels were placed on top of the gel with protein bands, followed by a thin layer of transparent film (Supplementary Fig. 1c, d) This configuration allows for a longer PISA observation of up to four hours.

A detailed description of the optical components and designs of PISA can be found in our previous report^29^. Briefly, PISA was built on a custom microscope body with two water-immersion objective lenses, fluorescence illumination (Special Optics, 54-10-7, NA = 0.66, 28.6×) and detection (Evident, XLUMPLFLN 20XW, NA = 1.0, 20×), that were placed below the coverslip at a tilted angle of 33.8 degrees. Bessel beam was generated by passing the laser source via an axicon lens (Mie Optics), which was then reflected by a Galvano mirror (Cambridge Technology, 6215HB), creating a light sheet. The detection port was connected to an EM-CCD camera (Andor, iXon Ultra 897) via an imaging lens and used to image fluorescence single molecules.

Imaging was conducted using a 647 nm fiber laser (MPB Communications, 2RU-VFL-P-2000-647) at 1,000 mW and detected through a near-infrared band-pass filter (Semrock, FF01-708/75-25). The protein bands embedded in polyacrylamide gel were imaged along a Y’-axis with a 4 μm step size at 50 ms exposure per frame. Images were acquired using a commercial software (Molecular Devices, MetaMorph) and a motorized stage (Prior Scientific, H117), which were controlled by a homemade program written in LabVIEW (National Instruments) and saved in TIFF format for further image analysis.

### Image analysis

All raw images were opened in ImageJ software (v1.51n), pre-processed using the rolling ball algorithm for background subtraction, and imported into Arivis Vision 4D (Zeiss) (Supplementary Fig. 5a, b). Denoising filter was applied to the background-subtracted images to improve the signal-to-noise ratio and a blob finder tool was used to detect spots in the 3D image matrix (Supplementary Fig.5c). Segment filter was optionally applied to remove single molecules with low-intensity signals. The images were saved and the acquired dataset containing the information of each spot, corresponding to single dye-labelled proteins, was exported in CSV format for data analysis.

## Data analysis

The exported dataset was analyzed and visualized using OriginPro (OriginLab) and Python (version 3.10.11). Frequency count function was used to generate a binned dataset and provide the count of dye-labelled proteins in each bin. The data was organized into a matrix of protein count × cell ID (rows × columns). To compensate for gel-to-gel variability, the proteome profile of each cell was manually aligned and scaled to standardize the peak (protein bands) positions across multiple technical or biological replicates. Molecular weights of unknown proteins were estimated by fitting a standard curve to relate the known molecular weights of proteins in a set of standards to their relative migration.

Two different normalization approaches were used to normalize the protein counts. For dimensional reduction such as principal component analysis (PCA), uniform manifold approximation and projection (UMAP), and diffusion map, the protein counts of individual cells were standardized using *z*-score, with the mean set to zero and standard deviation set to one. UMAP was used to visualize single cells in two (or three) dimensional maps and the clusters were identified using *k*-means.

Diffusion map was used to visualize the cell trajectory during cardiomyocyte differentiation by employing a random-walk algorithm to estimate the cell transition probabilities based on a weighted nearest neighbors graph^40^. Trajectory was inferred by computing diffusion pseudotime (DPT) for each cell relative to the root cell, and the pseudotemporal values were assigned to the cells along the inferred trajectory. To identify differentially regulated protein bands, the number of quantified proteins for each molecular weight was normalized by the total protein count in individual cells.

Differential protein bands (visualized in volcano plot) were determined by the Welch’s two-sided, two-sample *t*-test (*p* < 0.05, fold change > 1.2). For pairwise comparisons between three or more groups, significant differences were determined by the one-way Welch’s ANOVA with Games-Howell post-hoc test (*p* < 0.05).

The scRNA-seq datasets for D0, D15, and D30 were obtained from the ArrayExpress database at EMBL-EBI under the accession number E-MTAB-6268^41^. Three random samples (RS) were generated from the dataset, each containing 49 randomly selected single cells from D0, D15, and D30 (*n* = 12, 14, and 23, respectively). For a larger sample size, 3,000 single cells were randomly selected from D0, D15, and D30 (*n* = 1,000 for each sampling day). Differential genes were determined by the Welch’s two-sided, two-sample *t*-test (*p* < 0.05, fold change > 1.4). The analysis included a total of 10,484 genes, which represented all genes corresponding to proteins between 37 and 126 kDa. The protein molecular weights were obtained from UniProt.

## Data availability

Source Data are provided with this paper.

## Code availability

All custom analysis code generated as part of this work is available from the corresponding author upon reasonable request.

## Acknowledgements

We sincerely thank all members of our laboratory and Dr. Tomoyuki Ohkawa (Kyoto University) for the useful insights and discussions. L.K. acknowledges the MEXT scholarship program for its support. This work was supported by grants-in-aid for Scientific Research (A) (20H00460), Challenging Pioneering Research (19H05545 and 20K20458), Early-Career Scientists (19K15718 and 22K14800) from Japan Society for the Promotion of Science; and ACT-X (JPMJAX1914), PRESTO (JPMJPR25J4), and CREST (JPMJCR2334) from Japan Science and Technology Agency; and grants from the RIKEN DECODE project, Stage Transition project, RIKEN Incentive Research Projects, Suntory Rising Stars Encouragement Program in Life Sciences (SunRiSE). S.K. acknowledges support from RIKEN’s SPDR fellowship.

## Author contributions

Y.T., L.K., and S.K. conceived the main idea of the research. Y.T. designed the PISA microscope. S.K. and Y.T. constructed the microscopic system. L.K. and S.K. optimized the condition for microscopic imaging. L.K., S.K., T.H., and M.T. performed mammalian cell culture and cardiomyocyte differentiation from human induced pluripotent stem cells. L.K., Y.T., and S.K. wrote the manuscript, designed the experiments, and prepared the main and supplementary figures. L.K. designed the analysis pipelines and conducted the bioinformatical analysis. L.K. and M.T. performed all supplementary experiments. All authors contributed ideas for data analysis and interpretations and participated in the manuscript revision. Y.T. and S.K. supervised the research and acquired funding.

## Additional information

**Supplementary information** consists of 10 Supplementary figures, 2 Supplementary movies, and 1 Supplementary table.

## References

1. Papalexi, E. & Satija, R. Single-cell RNA sequencing to explore immune cell heterogeneity. Nat. Rev. Immunol. 18, 35–45 (2018).

2. Zhang, S., Li, X., Lin, J., Lin, Q. & Wong, K. C. Review of single-cell RNA-seq data clustering for cell-type identification and characterization. Rna 29, 517–530 (2023).

3. Jiang, L., Chen, H., Pinello, L. & Yuan, G. C. GiniClust: detecting rare cell types from single-cell gene expression data with Gini index. Genome Biol. 17, 1–13 (2016).

4. Xu, C. & Su, Z. Identification of cell types from single-cell transcriptomes using a novel clustering method. Bioinformatics 31, 1974–1980 (2015).

5. Taniguchi, Y. et al. Quantifying E. coli proteome and transcriptome with single-molecule sensitivity in single cells. Science 329, 533–538 (2010).

6. Schwanhäusser, B. et al. Global quantification of mammalian gene expression control. Nature 473, 337–342 (2011).

7. Darmanis, S. et al. Simultaneous multiplexed measurement of RNA and proteins in single cells. Cell Rep. 14, 380–389 (2016).

8. Gong, H. et al. Single-cell protein-mRNA correlation analysis enabled by multiplexed dual-analyte co-detection. Sci. Rep. 7, 2776 (2017).

9. Reimegård, J. et al. A combined approach for single-cell mRNA and intracellular protein expression analysis. *Commun*. Biol. 4, 624 (2021).

10. Vogel, C. & Marcotte, E. M. Insights into the regulation of protein abundance from proteomic and transcriptomic analyses. Nat. Rev. Genet. 13, 227–232 (2012).

11. Hanna, J., Guerra-Moreno, A., Ang, J. & Micoogullari, Y. Protein degradation and the pathologic basis of disease. Am. J. Pathol. 189, 94–103 (2019).

12. Kim, S., Kamarulzaman, L. & Taniguchi, Y. Recent methodological advances towards single-cell proteomics. Proc. Jpn. Acad. Ser. B 99, 306–327 (2023).

13. Mishra, N. C. Methodology for separation and identification of proteins and their interactions. Introduction to Proteomics 61–102 (John Wiley & Sons, Hoboken, 2010).

14. Wiśniewski, J. R., Hein, M. Y., Cox, J. & Mann, M. A “proteomic ruler” for protein copy number and concentration estimation without spike-in standards. Mol. Cell. Proteomics 13, 3497–3506 (2014).

15. Budnik, B., Levy, E., Harmange, G. & Slavov, N. SCoPE-MS: mass spectrometry of single mammalian cells quantifies proteome heterogeneity during cell differentiation. Genome Biol. 19, 1–12 (2018).

16. Zhu, Y. et al. Nanodroplet processing platform for deep and quantitative proteome profiling of 10–100 mammalian cells. Nat. Commun. 9, 882 (2018).

17. Li, Z. Y. et al. Nanoliter-scale oil-air-droplet chip-based single-cell proteomic analysis. Anal. Chem. 90, 5430–5438 (2018).

18. Woo, J. et al. High-throughput and high-efficiency sample preparation for single-cell proteomics using a nested nanowell chip. Nat. Commun. 12, 6246 (2021).

19. Petelski, A. A. et al. Multiplexed single-cell proteomics using SCoPE2. Nat. Protoc. 16, 5398–5425 (2021).

20. Gebreyesus, S. T. et al. Streamlined single-cell proteomics by an integrated microfluidic chip and data-independent acquisition mass spectrometry. Nat. Commun. 13, 37 (2022).

21. Leduc, A., Huffman, R. G., Cantlon, J., Khan, S. & Slavov, N. Exploring functional protein covariation across single cells using nPOP. Genome Biol. 23, 261 (2022).

22. Ctortecka, C. et al. An automated nanowell-array workflow for quantitative multiplexed single-cell proteomics sample preparation at high sensitivity. Mol. Cell. Proteomics 22, 100665 (2023).

23. Wang, Y. et al. Pick-up single-cell proteomic analysis for quantifying up to 3000 proteins in a mammalian cell. Nat. Commun. 15, 1279 (2024).

24. Ctortecka, C. et al. Automated single-cell proteomics providing sufficient proteome depth to study complex biology beyond cell type classifications. Nat. Commun. 15, 5707 (2024).

25. Bendall, S. C. et al. Single-cell mass cytometry of differential immune and drug responses across a human hematopoietic continuum. Science 332, 687–696 (2011).

26. Hughes, A. J., et al. Single-cell western blotting. Nat. Methods 11, 749–755 (2014).

27. Grist, S. M., Mourdoukoutas, A. P. & Herr, A. E. 3D projection electrophoresis for single-cell immunoblotting. Nat. Commun. 11, 6237 (2020).

28. Alibekova Long, M., Benman, W. K., Petrikas, N., Bugaj, L. J. & Hughes, A. J. Enhancing single-cell western blotting sensitivity using diffusive analyte blotting and antibody conjugate amplification. Anal. Chem. 95, 17894–17902 (2023).

29. Taniguchi, Y. et al. PISA: versatile microscope for 3D single molecule light sheet imaging. Preprint at 10.1101/2024.12.05.625331 (2024).

30. Patton, W. F. Detection technologies in proteome analysis. J. Chromatogr. B 771, 3–31 (2002).

31. Neuhoff, V., Arold, N., Taube, D. & Ehrhardt, W. Improved staining of proteins in polyacrylamide gels including isoelectric focusing gels with clear background at nanogram sensitivity using Coomassie Brilliant Blue G-250 and R-250. Electrophoresis 9, 255–262 (1988).

32. Jin, L. T., Hwang, S. Y., Yoo, G. S. & Choi, J. K. Sensitive silver staining of protein in sodium dodecyl sulfate-polyacrylamide gels using an azo dye, calconcarboxylic acid, as a silver-ion sensitizer. Electrophoresis 25, 2494–2500 (2004).

33. Chevallet, M., Luche, S. & Rabilloud, T. Silver staining of proteins in polyacrylamide gels. Nat. Protoc. 1, 1852–1858 (2006).

34. Bekker-Jensen, D. B. et al. An optimized shotgun strategy for the rapid generation of comprehensive human proteomes. Cell Syst. 4, 587–599 (2017).

35. Rajala, K., Pekkanen-Mattila, M. & Aalto-Setälä, K. Cardiac differentiation of pluripotent stem cells. Stem Cells Int. 2011, 383709 (2011).

36. Zakrzewski, W., Dobrzyński, M., Szymonowicz, M. & Rybak, Z. Stem cells: past, present, and future. Stem Cell Res. Ther. 10, 1–22 (2019).

37. Ivashchenko, C. Y. et al. Human-induced pluripotent stem cell-derived cardiomyocytes exhibit temporal changes in phenotype. Am. J. Physiol. Heart Circ. Physiol. 305, H913–H922 (2013).

38. Hellen, N. et al. Proteomic analysis reveals temporal changes in protein expression in human induced pluripotent stem cell-derived cardiomyocytes in vitro. Stem Cells Dev. 28, 565–578 (2019).

39. Jabart, E. et al. Single-cell protein expression of hiPSC-derived cardiomyocytes using single-cell westerns. J. Mol. Cell. Cardiol. 149, 115–122 (2020).

40. Haghverdi, L., Büttner, M., Wolf, F. A., Buettner, F. & Theis, F. J. Diffusion pseudotime robustly reconstructs lineage branching. Nat. Methods 13, 845–848 (2016).

41. Friedman, C. E. et al. Single-cell transcriptomic analysis of cardiac differentiation from human PSCs reveals HOPX-dependent cardiomyocyte maturation. Cell Stem Cell 23, 586–598 (2018).

42. Doll, S. et al. Region and cell-type resolved quantitative proteomic map of the human heart. Nat. Commun. 8, 1469 (2017).

43. Liu, Y. et al. Robustness of single-cell RNA-seq for identifying differentially expressed genes. BMC Genomics 24, 371 (2023).

44. Narsinh, K. H. et al. Single-cell transcriptional profiling reveals heterogeneity of human induced pluripotent stem cells. J. Clin. Invest. 121, 1217–1221 (2011).

45. Nguyen, Q. H. et al. Single-cell RNA-seq of human induced pluripotent stem cells reveals cellular heterogeneity and cell state transitions between subpopulations. Genome Res. 28, 1053–1066 (2018).

46. Shao, W. et al. Comparative analysis of mRNA and protein degradation in prostate tissues indicates high stability of proteins. Nat. Commun. 10, 2524 (2019).

47. Podgorski, K., Terpetschnig, E., Klochko, O. P., Obukhova, O. M. & Haas, K. Ultra-bright and-stable red and near-infrared squaraine fluorophores for in vivo two-photon imaging. PLoS One 7, e51980 (2012).

48. Liu, X., Jiang, Y., Cui, Y., Yuan, J. & Fang, X. Deep learning in single-molecule imaging and analysis: recent advances and prospects. Chem. Sci. 13, 11964–11980 (2022).

49. Savas P. et al. Single-cell profiling of breast cancer T cells reveals a tissue-resident memory subset associated with improved prognosis. Nat. Med. 24, 986–993 (2018).

50. Slavov N. Single-cell protein analysis by mass spectrometry. Curr. Opin. Chem. Biol. 60, 1–9 (2021).

51. Kinoshita, E., Kinoshita-Kikuta, E. & Koike, T. Separation and detection of large phosphoproteins using Phos-tag SDS-PAGE. Nat. Protoc. 4, 1513–1521 (2009).

52. Gough, P. & Myles, I. A. Tumor necrosis factor receptors: pleiotropic signaling complexes and their differential effects. Front. Immunol. 11, 585880 (2020).

53. Jin, J. et al. Robotic data acquisition with deep learning enables cell image–based prediction of transcriptomic phenotypes. Proc. Natl Acad. Sci. USA 120, e2210283120 (2023).

54. LI-COR Biosciences. In-Gel Westerns. LI-COR. https://www.licor.com/bio/applications/in-gel-westerns (accessed December 10, 2024).

55. Burridge, P. W. et al. Chemically defined generation of human cardiomyocytes. Nat. Methods 11, 855–860 (2014).

