## Supplementary Information for "Comprehensive molecular profiling of single-cell proteome via gel electrophoresis and 3D single-molecule imaging"

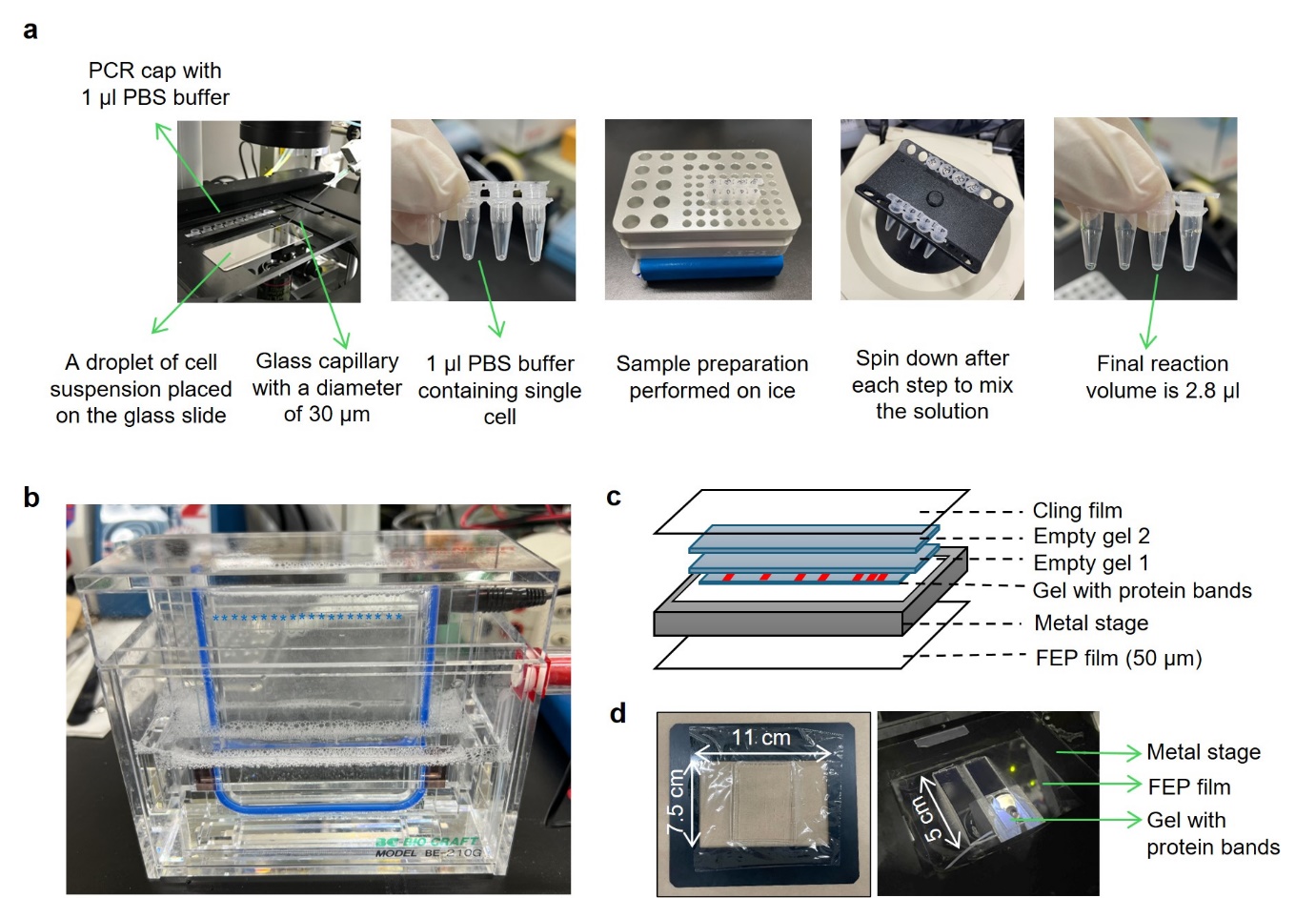
**Supplementary Figure 1 | Workflow of single-cell PAGE-PISA.** **a** Images of manual single-cell isolation and microliter-scale single-cell sample preparation in PCR tubes. Briefly, the single cells were manually collected using the 30 µm G-tip low-binding coated micro-glass needle, transferred to the caps of the PCR tubes containing 1 µL of PBS buffer, and spun down. All sample preparation steps were performed on ice and mixed by spinning down. The final reaction volume after adding lysis buffer, protease inhibitor, dye, and quencher solution is 2.8 µL. **b** Image of single-cell electrophoresis. Asterisks represent the number of loading wells. Each loading well has a 4 mm width and can load up to 8 µl sample volume. **c, d** Schematic drawing (**c**) and images (**d**) of single-cell PAGE-PISA setup before PISA imaging. The FEP film was taped to the metal stage with an open window area of 11 cm × 7.5 cm and irradiated under UV for one hour to remove autofluorescence dust. After electrophoresis, the polyacrylamide gel was cut vertically into a 5 cm length, approximately from 20‒40 to 100‒150 kDa, and placed on the FEP film. Two unused polyacrylamide gels (empty gels) were used to cover the gel containing protein bands, followed by a thin layer of transparent film. The setup was then placed on a microscopic stage for PISA imaging (right).

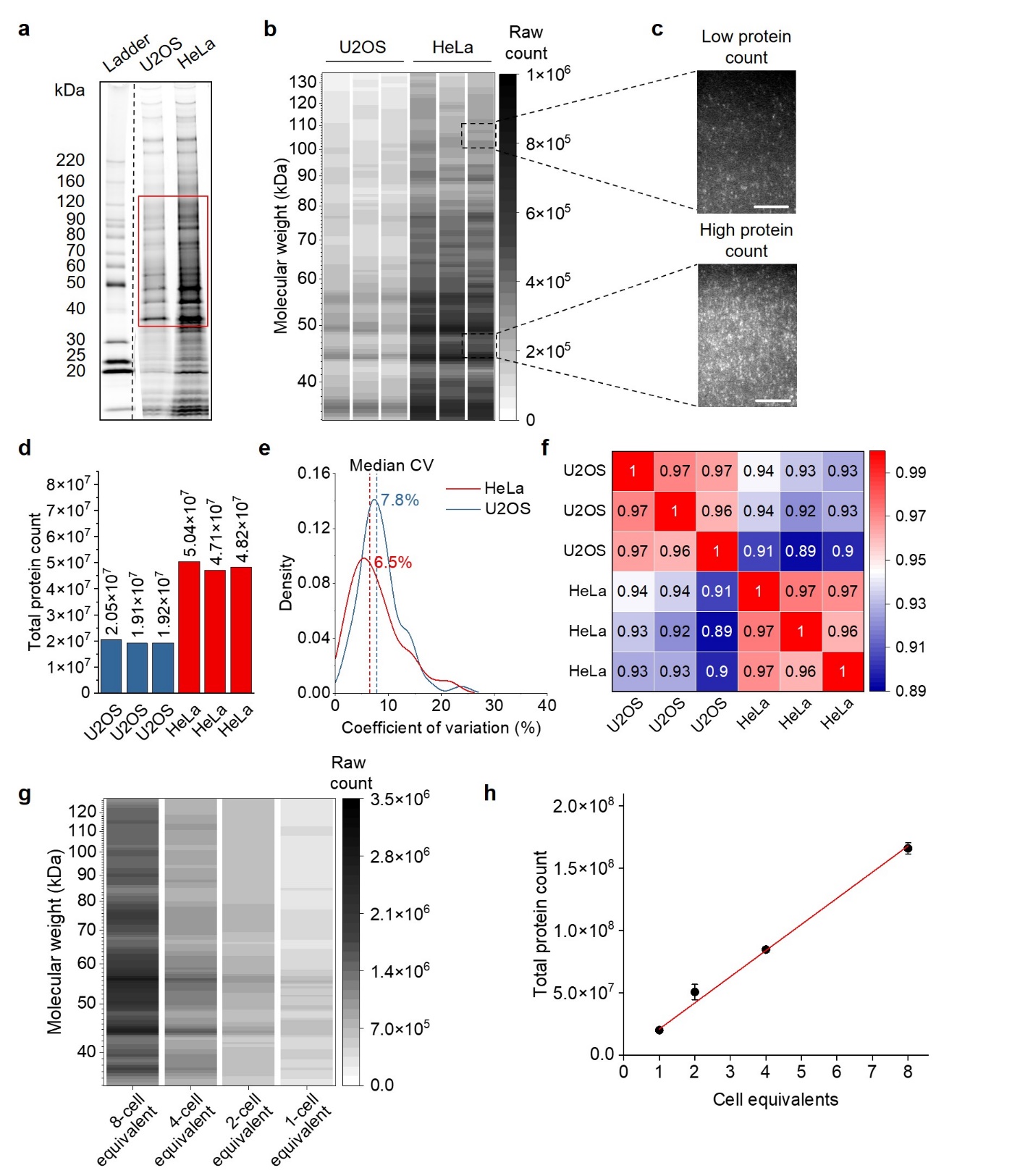
**Supplementary Figure 2 | Single-molecule imaging of highly diluted bulk cell lysates.** **a** SDS-PAGE gel image of dye-labelled proteins from approximately 7.5 × 10^3^ U2OS and HeLa cell lysates visualized by the standard gel imager. Samples and molecular weight ladder were run on the same gel and the image was cropped (indicated by a dotted line) only for the purpose of this figure. **b** Electropherogram of the bulk cell lysates diluted to an approximately single-cell level obtained by single-molecule imaging using PISA. The electropherogram was reconstructed based on the raw protein counts from 35‒135 kDa, as indicated by the red box in panel (**a**). **c** Representative raw images of the single molecules during PISA imaging, shown at two molecular weight regions with different protein abundances. Each detected single molecule represents a single dye-labelled protein. Scale bar: 50 µm. **d** Total protein count of individual replicates of bulk HeLa and U2OS cell lysates diluted to single-cell level. **e** Distribution of coefficient of variation (CV) of protein counts across molecular weights, showing medians of 7.8% and 6.5% for U2OS and HeLa, respectively. **f** Reproducibility assessment of the protein counts across three technical replicates for each cell type. **g** Electropherogram of bulk U2OS cell lysates diluted to 1, 2, 4, and 8 cells equivalent, shown across 35‒128 kDa. Each electropherogram represents the average raw protein counts across triplicates. **h** Linearity between cell equivalents and total protein count. Error bars represent the standard error of the mean. Number of technical replicates: *n* = 3 for U2OS and *n* = 3 for HeLa cells.

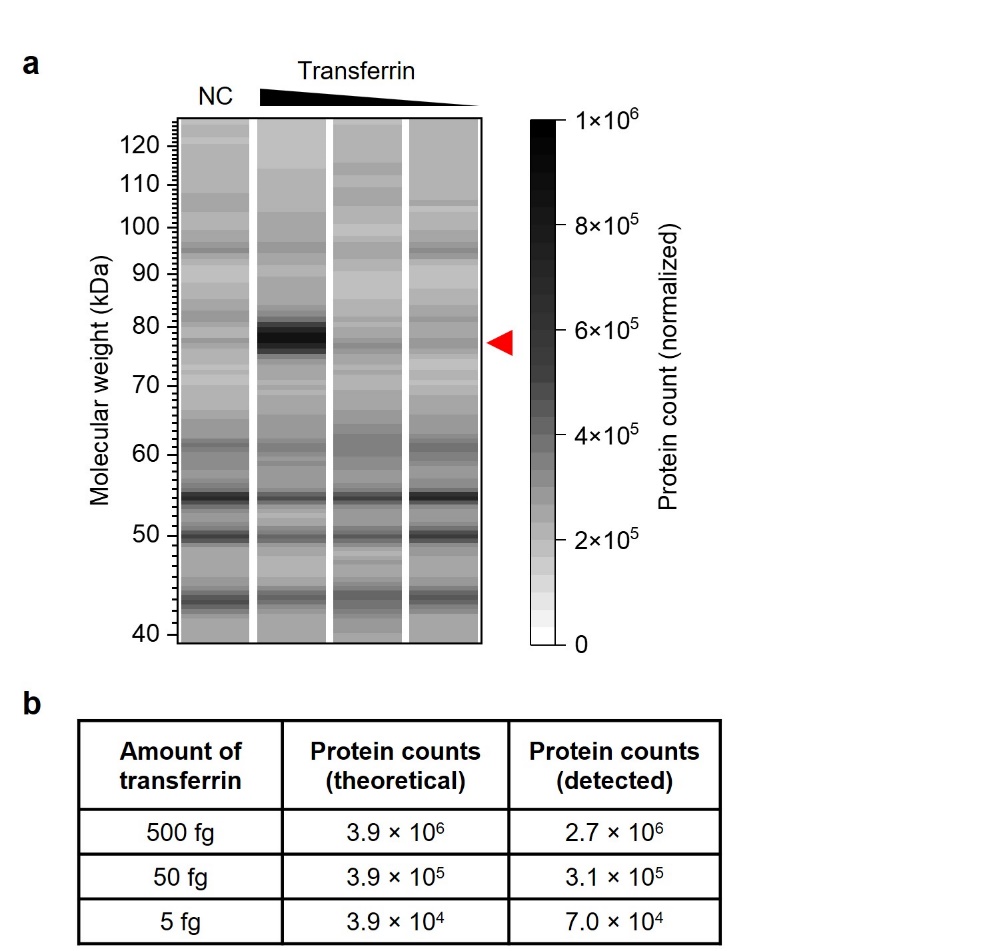
**Supplementary Figure 3 | Detection of low-abundance protein by single-cell PAGE-PISA using cell lysate samples spiked with purified transferrin. a** Electropherogram of lysate spike-in samples, shown across 39‒128 kDa. The lanes represent negative control (NC) and cell lysates spiked with 500 fg, 50 fg, and 5 fg transferrin. Each electropherogram represents the average protein counts across triplicates, normalized to control. The red arrow indicates the transferrin protein band at 77 kDa. **b** Quantitative comparison between theoretical and average protein counts of transferrin detected after subtraction of the negative control. Number of technical replicates: *n* = 3 per condition.

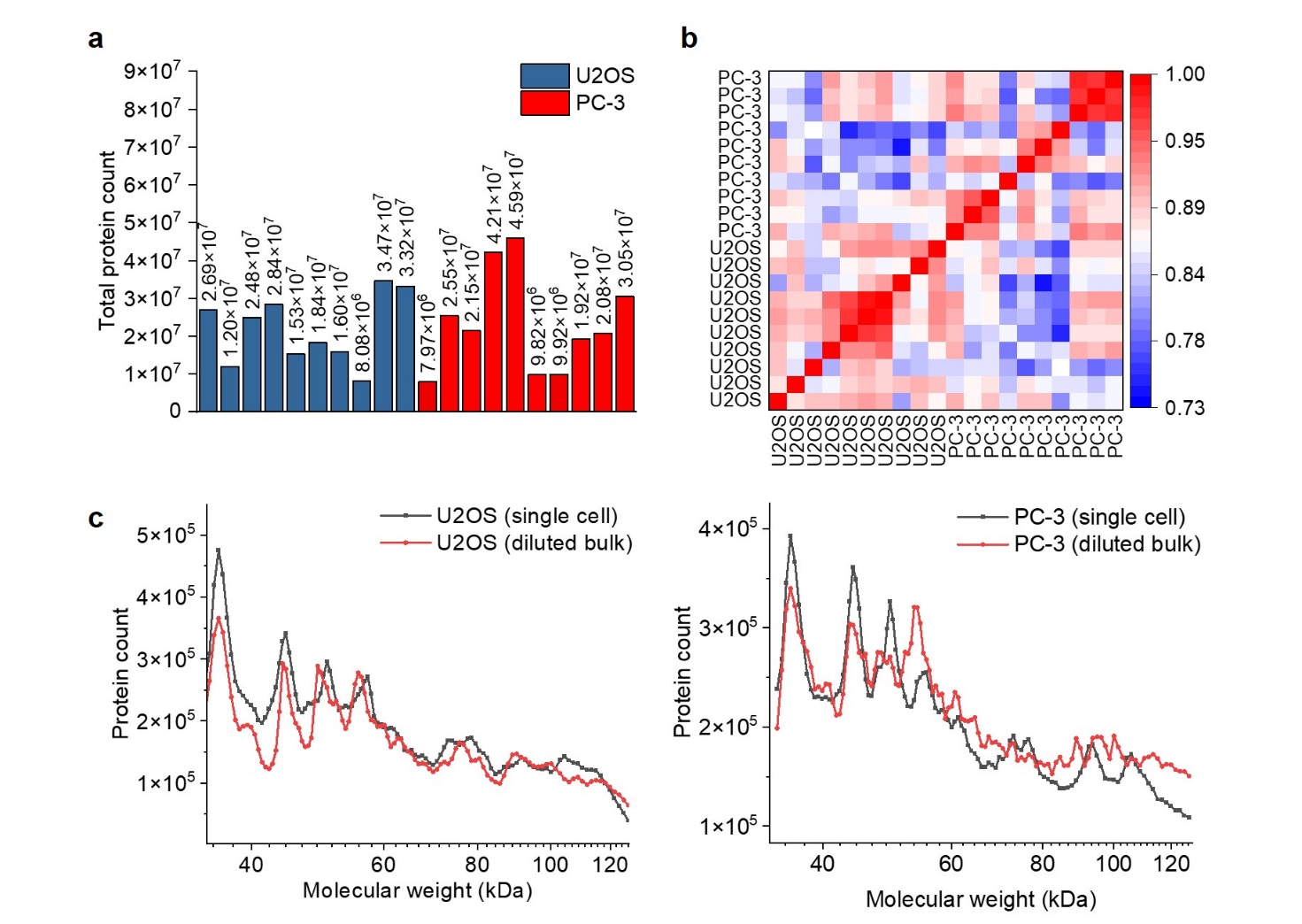
**Supplementary Figure 4 | Comparative analysis between single cells and highly diluted bulk cell lysates equivalent to single-cell level. a** Total protein count of individual U2OS and PC-3 cells from 35‒128 kDa. **b** Pearson correlation across 10 biological replicates for each cell type, showing the reproducibility of protein counts across molecular weights. **c** Average proteome profiles across molecular weights for U2OS (left) and PC-3 (right) cells, comparing single-cell lysates (black line) with bulk cell lysates diluted to single-cell level (red line). The proteome profiles were averaged independently across multiple single-cell lysates and across technical replicates of bulk cell lysates diluted to single-cell level, respectively. Number of biological replicates (single cells): *n* = 10 for U2OS and *n* = 10 for PC-3, number of technical replicates (bulk cell lysates): *n* = 3 for U2OS and *n* = 3 for PC-3 cells.

**Supplementary Figure 5 | Detection of single molecules. a, b** Images taken during PISA measurement showing raw (**a**) and background-subtracted (**b**) images of single-cell lysate (left) and negative control (right). Images were taken under the same microscopic condition and the intensity was normalized between the raw and negative control images. Each detected single molecule represents a single dye-labelled protein. **c** Image analysis for single-molecule detection. After background subtraction, the image was processed using discrete Gaussian filter (left). Single molecules were identified using the blob finder tool, highlighted as blue pixels (center). Segment filter was subsequently applied to remove single molecules with low-intensity signals that were indistinguishable from the background (right). Yellow circles indicate representative single molecules that were filtered out after threshold application. Scale bar: 50 µm.
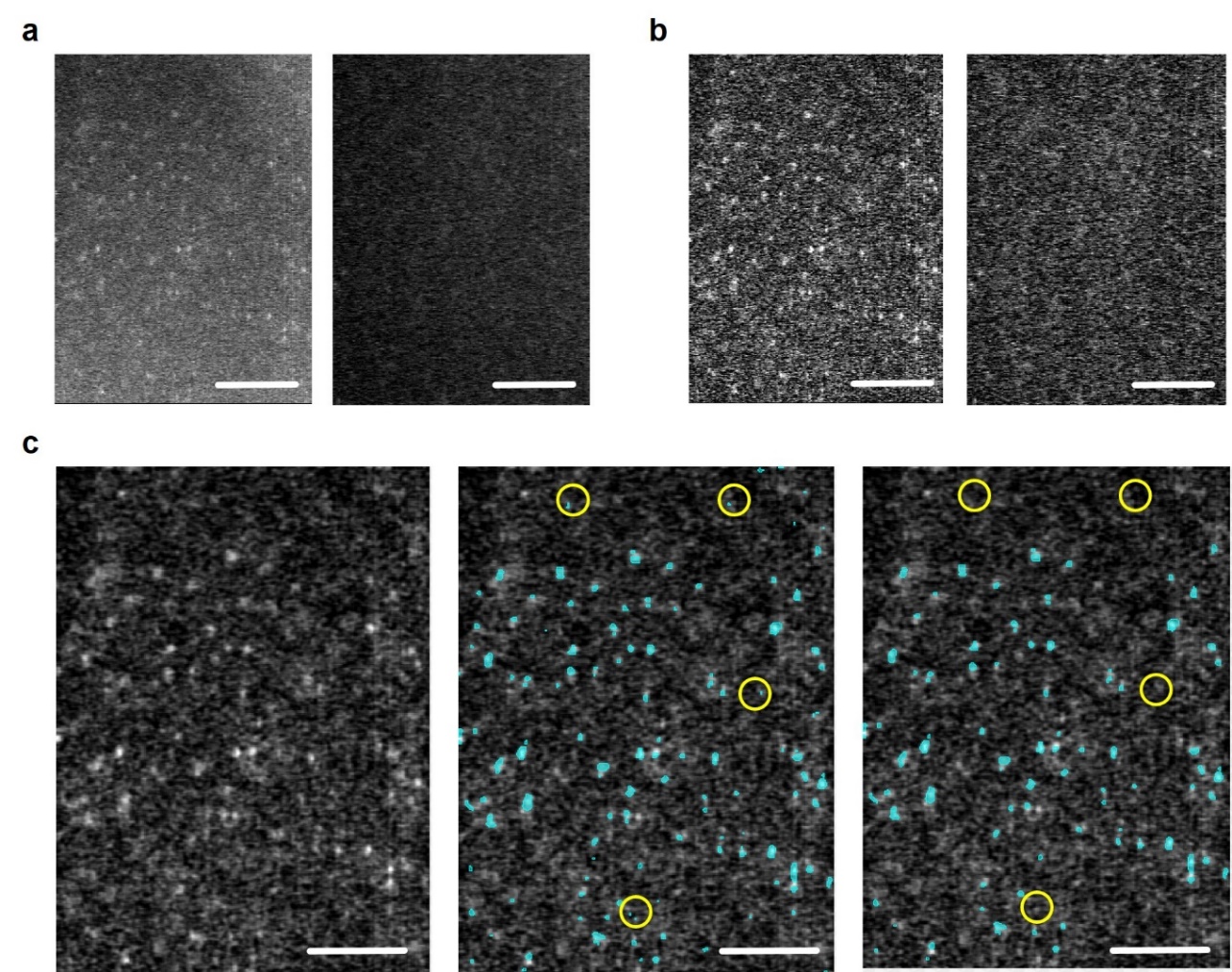

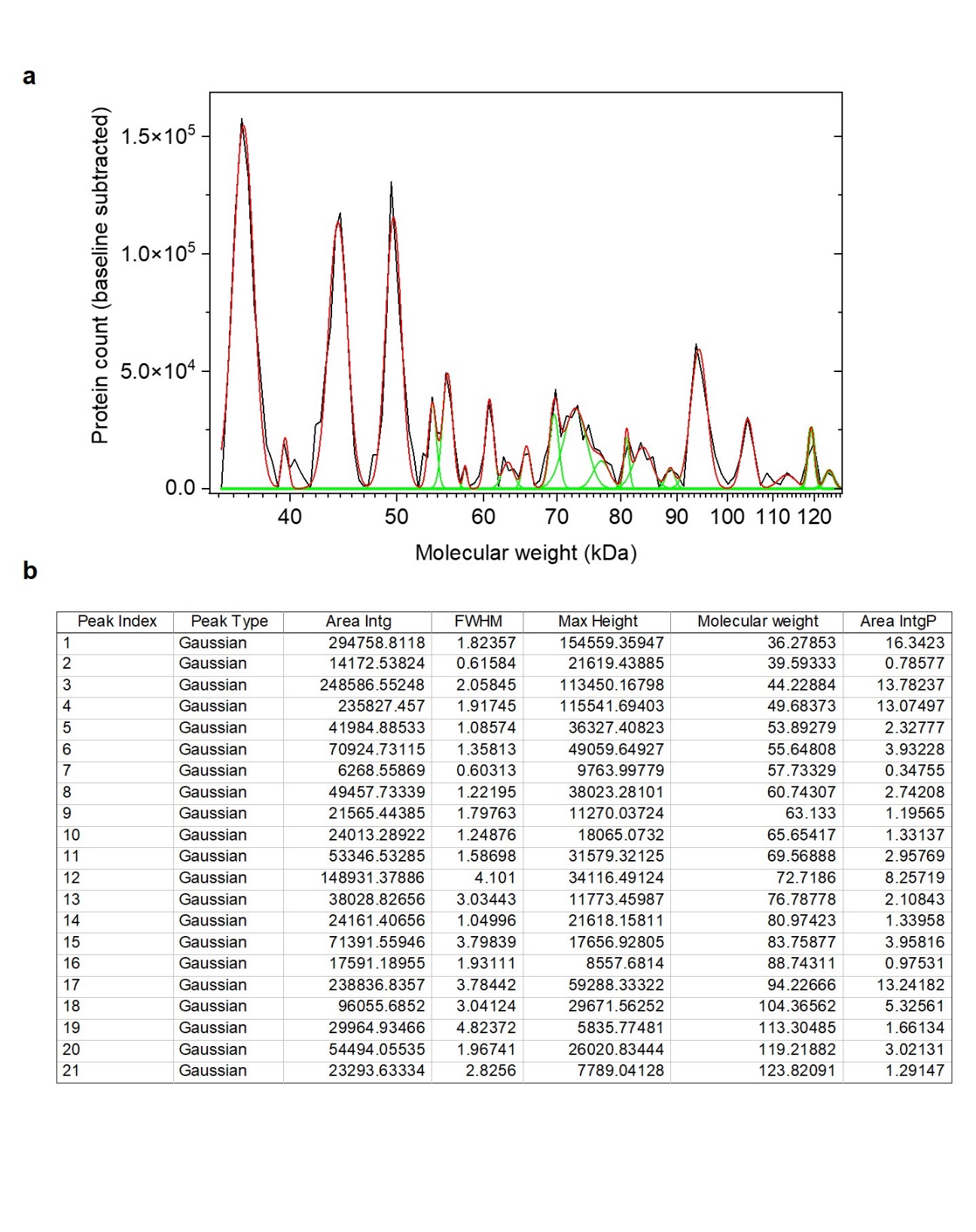
**Supplementary Figure 6 | Resolution power analysis of a representative separation profile from a single-cell lysate. a** Peak fitting of the single-cell proteome profile across 35–128 kDa range. The proteome profile (black line) is overlaid with the fitted model (red line), composed of 21 Gaussian peaks (green curves). Peaks were identified using second derivative-based detection following baseline subtraction. **b** Table summarizing the fitted parameters for each peak, including peak area, position, height, and full width at half maximum (FWHM). Peak fitting was performed using constrained nonlinear regression with the following bounds: (1) peak position ±5%, (2) FWHM ±50%, and (3) peak height ±50% of the initial estimates generated by OriginPro.

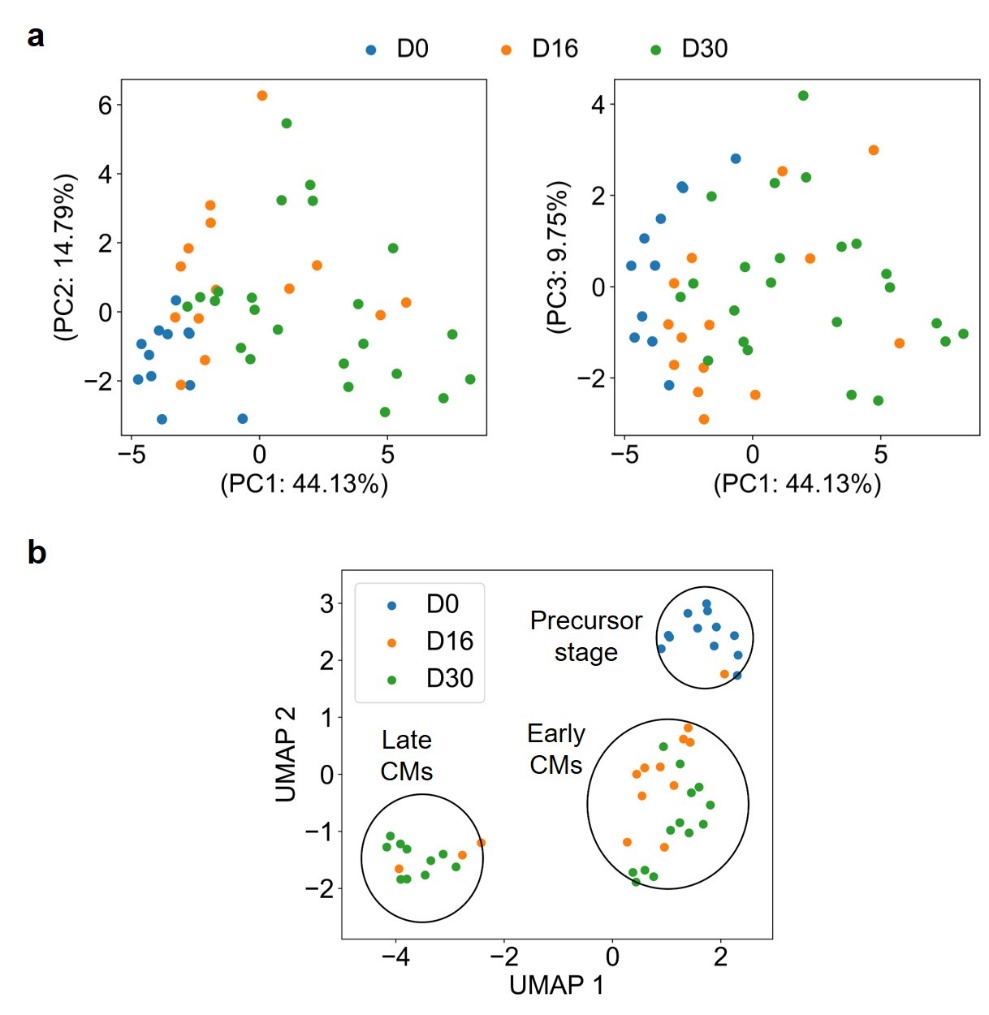
**Supplementary Figure 7 | Dimensionality reduction analysis of single hiPSCs and hiPSC-CMs obtained by single-cell PAGE-PISA. a** Unsupervised principal component analysis (PCA) of single cells (hiPSCs: D0, hiPSC-CMs: D16 and D30) based on the standardized protein counts from 35‒112 kDa. The single cells coloured by the sampling days were separated in a time-wise manner along the first principal component (PC1). Number of biological replicates: *n* = 12 for D0, *n* = 14 for D16, and *n* = 23 for D30. **b** Uniform manifold approximation and projection (UMAP) analysis with *k*-means clustering of the single cells. Three clusters were identified using *k*-means, representing different developmental stages during cardiomyocyte differentiation. Number of cells in each cluster: *n* = 13 for the precursor stage, *n* = 23 for early cardiomyocytes, and *n* = 13 for late cardiomyocytes.

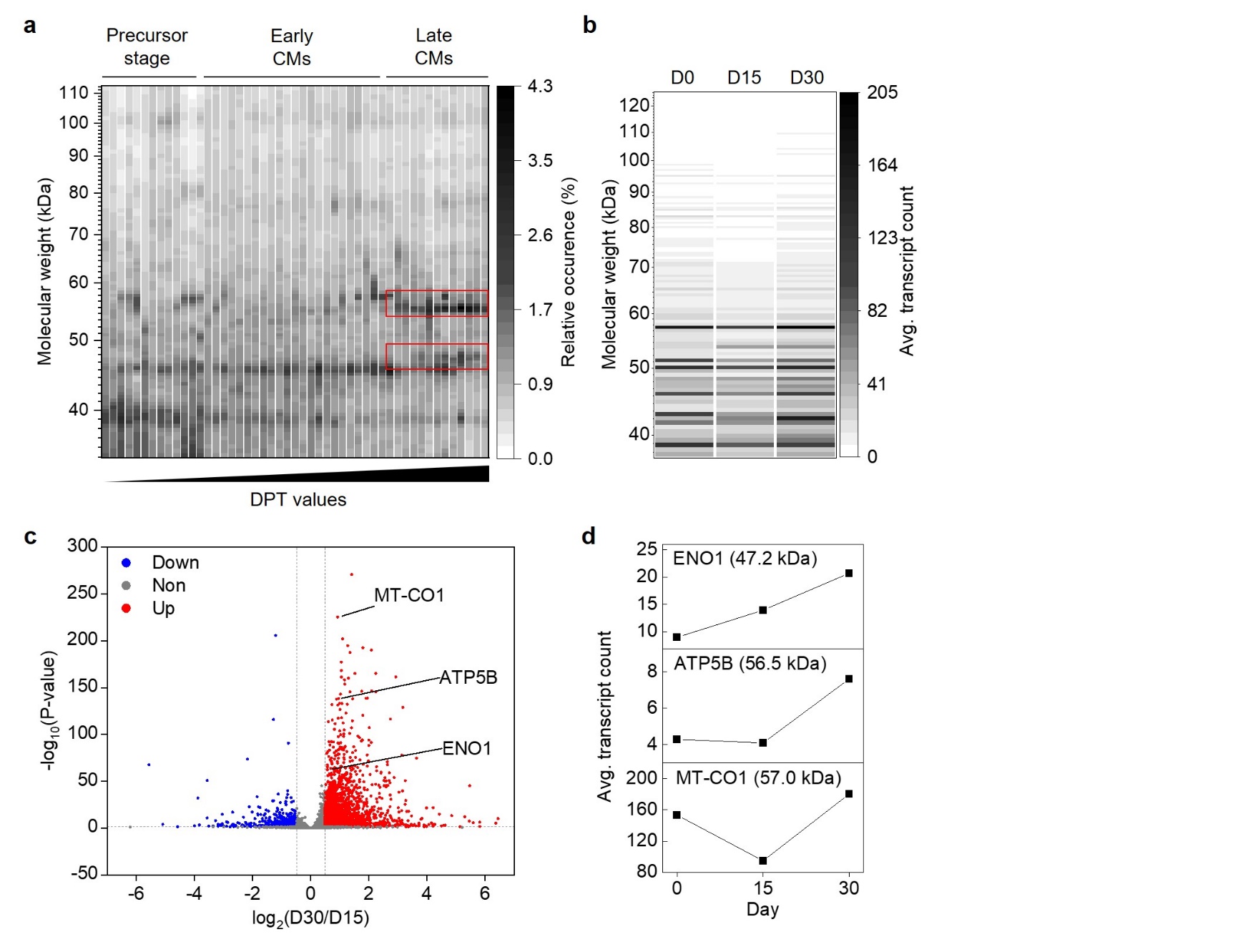
**Supplementary Figure 8 | Single-cell proteome and transcriptome analysis during cardiomyocyte differentiation. a** Reconstructed single-cell electropherogram of the precursor stage (*n* = 13), early cardiomyocytes (*n* = 23), and late cardiomyocytes (*n* = 13). The electropherogram was arranged based on ascending DPT values of single cells, from left to right, and was reconstructed based on the normalized protein counts from 35‒112 kDa. Differentially expressed protein bands at 55.0‒57.7 kDa and 47.1‒48.0 kDa in late cardiomyocytes were indicated by the red box. **b** Heatmap of the scRNA-seq data showing the average transcriptome profiles of D0, D15, and D30. **c** Volcano plot showing differentially regulated genes in comparative analysis between D30 and D15. Differential genes were determined by Welch's two-sided, two-sample *t*-test (*p* < 0.05, fold change > 1.4). **d** Average transcript counts of D0, D15, and D30 cells showing temporal gene expression of *ENO1*, *ATP5B*, and *MT-CO1* during cardiomyocyte differentiation. In (**b**, **c**), only genes with corresponding proteins from 37‒126 kDa were included in the analysis. In (**b**, **d**), the transcript counts were averaged across multiple single cells: D0 (*n* = 9,146), D15 (*n* = 2,897), and D30 (*n* = 3,294).

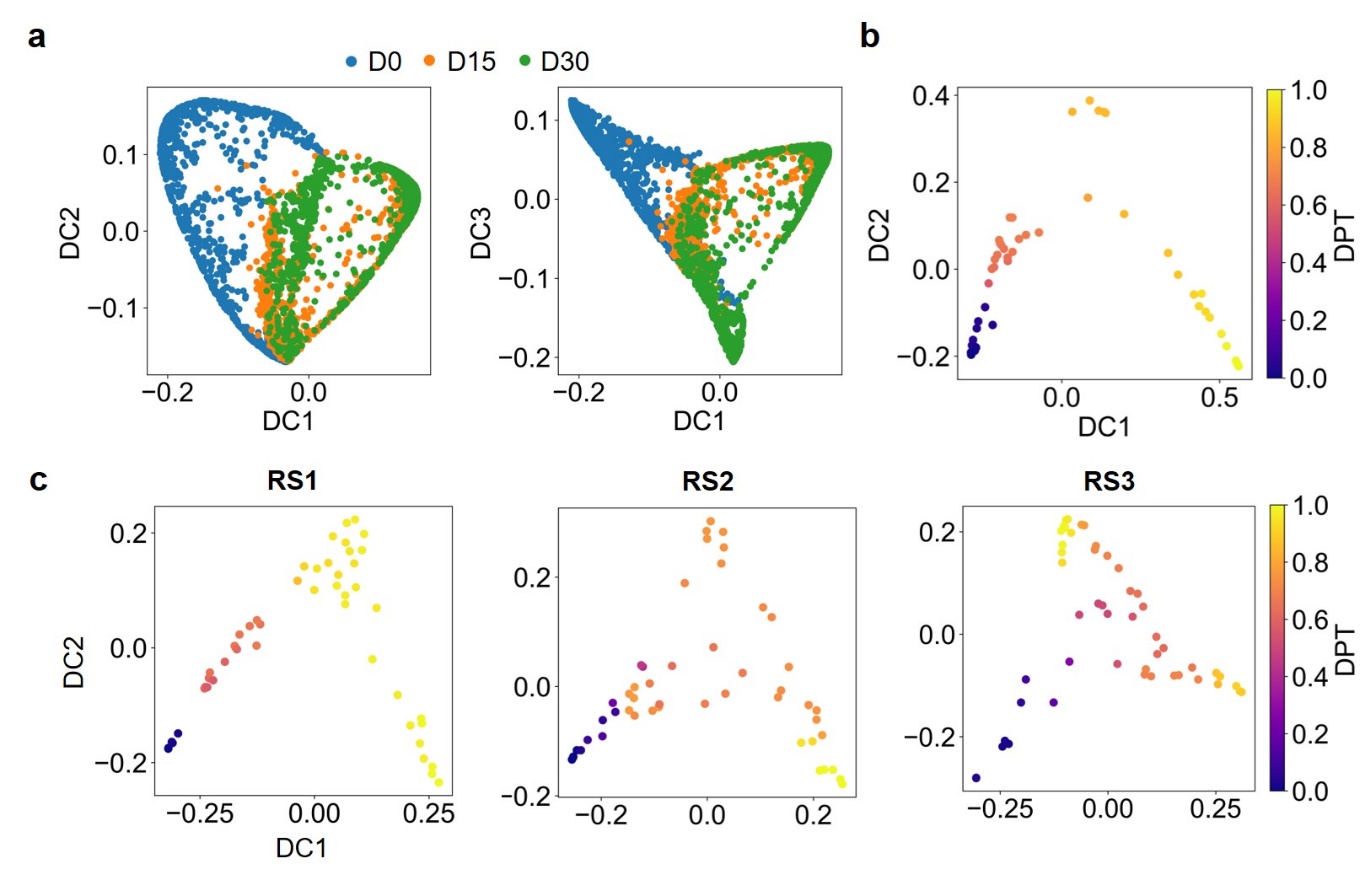
**Supplementary Figure 9 | Diffusion pseudotime reveals the temporal ordering of single cells during cardiomyocyte differentiation. a** Diffusion maps showing single-cell transcriptome data arranged along a pseudotemporal trajectory based on the transition probabilities between two cells. The single cells were coloured with respect to their sampling days. Number of biological replicates: *n* = 1,000 each for D0, D15, and D30. **b, c** Cell trajectories based on diffusion pseudotime for single-cell proteome (**b**) and transcriptome data (**c**). The trajectory starts from the root cell (blue) that is located at the left terminal and progresses towards the most differentiated cells that are located furthest from the root (yellow). DC refers to diffusion components, DPT refers to diffusion pseudotime, and RS refers to random sample. Number of biological replicates for single-cell PAGE-PISA: *n* = 12, 14, and 23 for D0, D16, and D30, respectively. Number of biological replicates for scRNA-seq: *n* = 12, 14, and 23 for D0, D15, and D30, respectively.

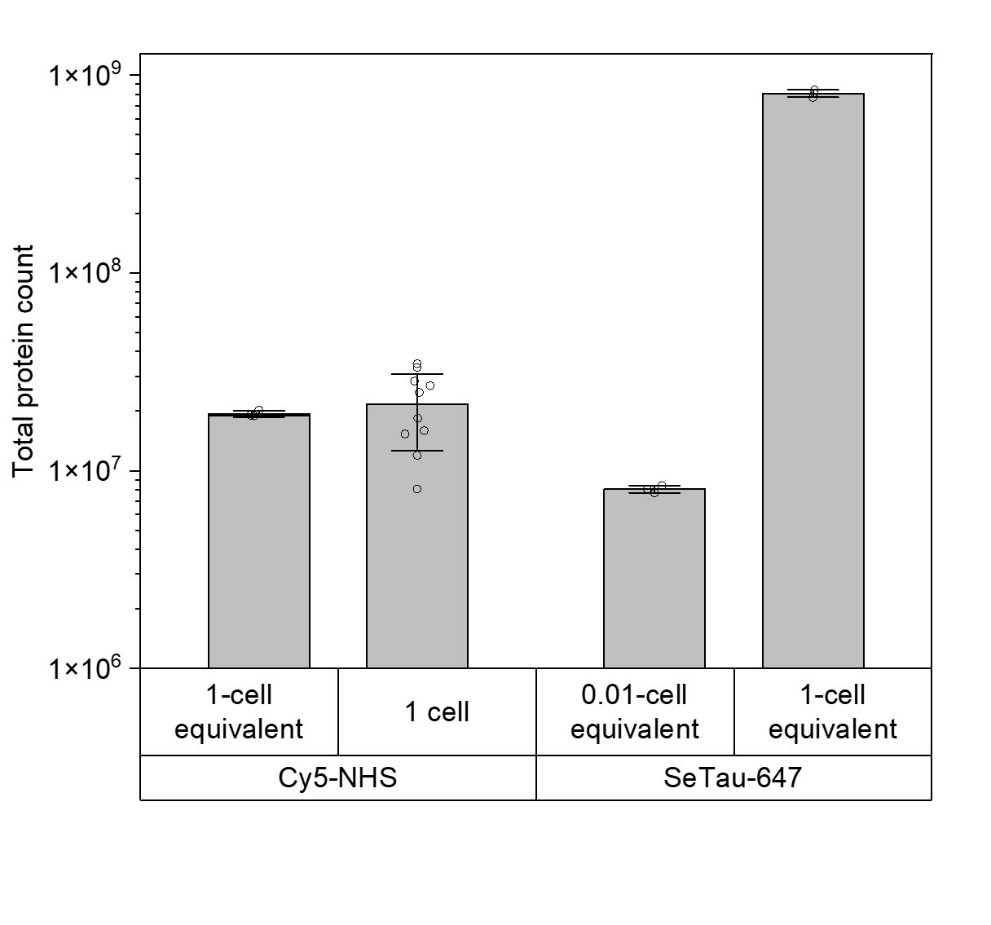
**Supplementary Figure 10 |** Comparison of total protein count using Cy5 and SeTau fluorescent dyes. Cy5-NHS ester dye was used to label either single-cell lysates (from individual cells) or bulk cell lysates diluted to 1-cell equivalent. SeTau-647-NHS ester dye was used to label bulk cell lysates at 1-cell or 0.01-cell equivalent. SeTau-labelled 1-cell equivalent cell lysates showed a 1–2 order of magnitude increase in total protein counts compared to Cy5-labelled cell lysates. Error bars represent ± standard deviation. Number of biological replicates (single cells): *n* = 10, number of technical replicates (bulk cell lysates): *n* = 3.

**Supplementary Table 1. Comparison between single-cell PAGE-PISA and MS-based single-cell proteomics approaches**

| Feature | Single-cell PAGE-PISA | MS-based single-cell proteomics |
| --- | --- | --- |
| Quantification | Provides absolute molecular counts via single-molecule fluorescence imaging. Quantifies approx. 1% of the total protein abundance. | Provides relative abundance via label-free or isotopic labelling. |
| Sensitivity | Detects 10^4^–10^5^ copies per protein species. | Detects 10^4^–10^5^ copies per protein species. |
| Protein identification | Currently, 20–60 protein bands can be resolved simultaneously via molecular weight. No direct protein identification but possible with antibody-based detection directly on gel or in combination with FACS and conventional MS. | Identifies >3, 000 protein groups via mass spectra and database matching. |
| Throughput | 20 cells per experiment but can be scaled up to ~1,000 cells through automation within 30 hours. | High-throughput processing of 100–1,000 cells within several days. |
| Cost & workflow | Cost-effective and straightforward ($0.6–1/cell) | Expensive and technically complex ($2–3/cell) |
| Applications | Whole proteome profiling, analysis of intact, native, modified, and spatially confined proteins | Proteome-wide identification, de novo discovery, spatial proteomics through imaging |
